# The gene regulatory basis of genetic compensation during neural crest induction

**DOI:** 10.1101/314534

**Authors:** Christopher M. Dooley, Neha Wali, Ian M. Sealy, Richard J. White, Derek L. Stemple, John E. Collins, Elisabeth M. Busch-Nentwich

## Abstract

**Background:** The neural crest (NC) is a vertebrate-specific cell type that contributes to a wide range of different tissues across all three germ layers. The gene regulatory network (GRN) responsible for the formation of neural crest is conserved across vertebrates. Central to the induction of the NC GRN are *AP-2* and *SoxE* transcription factors but detailed interactions within the network remain to be resolved.

**Results:** We have used gene knockout and RNA sequencing strategies to dissect NC differentiation in zebrafish. We establish that initiation of the NC GRN takes place just after genome activation. We genetically ablate the NC using double mutants of *tfap2a;tfap2c* or remove specific subsets of the NC with *sox10* and *mitfa* knockouts and characterise genome-wide gene expression levels across multiple time points. We find that although a single allele of *tfap2c* is capable of maintaining early NC induction and differentiation in the absence of *tfap2a* function, expression of many target genes remains abnormal and sensitive to *tfap2* dosage. This separation of morphological and molecular phenotypes identifies a core set of genes required for early NC development. Using gene knockouts, we associate previously uncharacterised genes with pigment cell development and establish a role for maternal Hippo signalling in melanocyte differentiation.

**Conclusions:** Stepwise genetic ablation of the NC identifies the core gene module required for neural crest induction. This work extends and refines the NC GRN while also uncovering the complex transcriptional basis of genetic compensation via paralogues.

## Background

Development from a single fertilised cell to the complex adult form requires a simultaneously robust and plastic gene regulatory program. The neural crest is a transient pluripotent stem cell population capable of crossing germ layer boundaries and differentiating into highly diverse tissue types while migrating long distances in the developing embryo. The establishment of the neural crest and its subsequent tissue derivatives is specific to vertebrates and has played a fundamental role in their variation and evolutionary success [1–3]. Neural crest cells require a complex combination of external inductive signals such as Wnts, Fgfs, Notch/delta and Bmps (Fig. 1a). These extrinsic signals can be considered the first phase of the neural crest gene regulatory network (GRN) followed by a second phase of tightly controlled intrinsic gene expression. Two of these intrinsic signals of fundamental importance for evolution and development of the neural crest that set vertebrates apart from other chordates such as amphioxus and tunicates are the *AP-2* and *SoxE* genes families. [4–8].

**Figure 1.**
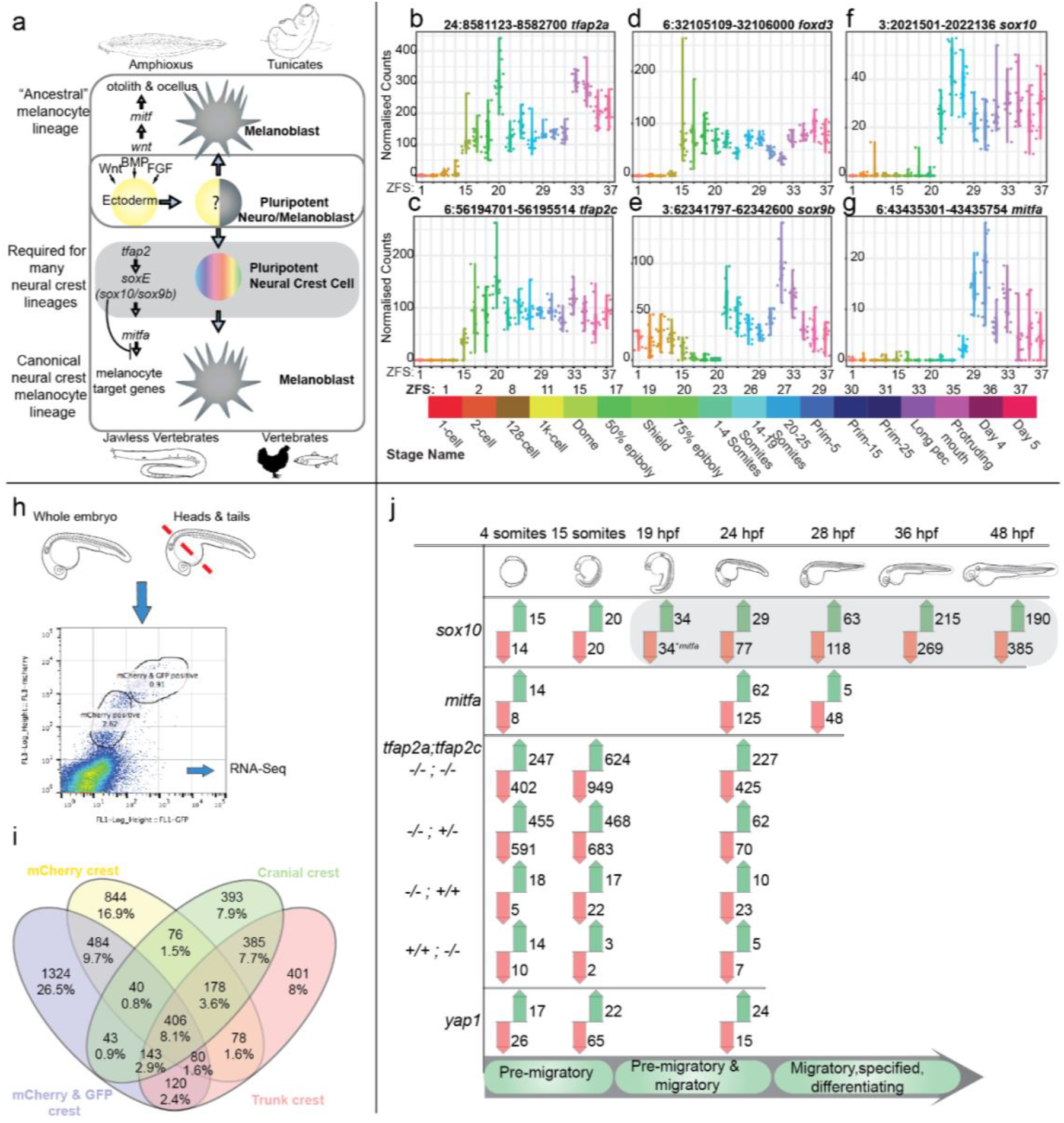
Analysis of the zebrafish NC GRN using gene expression data, knockouts and tissue-specific sequencing. **a** The NC is induced by different morphogens, for example Wnt, BMP and FGF acting on ectoderm. Non-vertebrate chordates lack NC cells but are capable of producing pigmented cells and otoliths via *mitf*. AP-2 and SoxE family genes are required in vertebrates to form the NC and these also contribute to the differentiation of specific NC tissues types. **b-g** 3’ end transcriptome sequencing (DeTCT) of six key neural crest transcription factors (*tfap2a*, *tfap2c*, *foxd3*, *sox9b*, *sox10*, *mitfa*) across 18 developmental time points covering zygote to 5 dpf. Normalised counts of individual embryos are plotted for each stage. The mapped Zv9 genomic positions of each 3’end are at the top of the plots next to the gene names. ZFS numbers are labelled with their corresponding stage names and representative colouring. **h** FACS of dissociated *sox10:mg* was sorted based on mCherry and GFP signals at 22-23 hpf and were either sorted as whole embryos or separated heads and tails. Multiple replicates of each cell population were harvested and sequenced via RNA-Seq. **i** FACS transgenic populations were compared to non-transgenic populations using DESeq2 to produce gene enrichment lists for each population. The enriched gene lists for the mCherry and mCherry/ GFP population from whole embryos and mCherry and/or GFP positive populations from the head or trunk were then compared to each other as a Venn diagram. **j** An overview of the transcriptomics loss of function analysis using 3’ tag sequencing, carried out at stages of premigratory, migratory and differentiating neural crest cells. The phases of NC differentiation are noted at the bottom. Changing genes at adj. p-value <0.05 when compared to wild-type siblings are represented with green arrows for increased and with red arrows for decreased abundance. The *sox10* downstream target *mitfa* is first detected as reduced at 19 hpf.

Mutations in neural crest genes lead to disease in humans, highlighting the importance of this cell population for human health. Animal models faithfully recapitulate these defects demonstrating functional conservation. In humans and mice, mutations in *TFAP2A* lead to branchio-oculo-facial syndrome presenting as defects in cranial development and cranial closure [9,10]. Similarly, mutations in zebrafish *tfap2a* lead to craniofacial defects in addition to a reduction in melanocytes [11,12]. The Tfap2 family arose from a single gene in a chordate ancestor that underwent gene duplication resulting in five family members in zebrafish. Removing combinations of *tfap2* family membersresults in a wide array of phenotypes. For example, the neural crest is completely ablated in *tfap2a*;*tfap2c* zebrafish whereas there is a dramatic and specific reduction of melanocytes in *tfap2a*;*tfap2e* zebrafish embryos [13–18]. Furthermore, melanomas, squamous cell carcinomas, most skin and breast cancers and a few cervical and urothelial cancers have strong nuclear immunoreactivity for TFAP2A. [19,20].

Haploinsufficiency of *SOX10* results in Waardenburg syndrome; patients exhibit defects in the peripheral and enteric nervous systems and also pigmentation defects [21,22]. Similar to humans, mice affected by the Dominant megacolon mutation *Sox10^Dom^* also have defects in melanocyte development, enteric neuron defects and develop megacolon as heterozygotes [23]. Homozygous knockouts of murine *Sox10* are embryonic lethal and also lead to strong myelination phenotypes and an overall lack of peripheral glia [24].

The expression of *sox10* is first detectable in premigratory neural crest cells and expression is maintained in certain neural crest linages, for example glia, but reduced in many other neural crest-derived tissues in zebrafish [25–27] and mouse [28–30]. Following neural crest induction, *sox10* plays a vital role in the establishment of non-ectomesenchymal neural crest cells in zebrafish. Knockouts in zebrafish *sox10* behave in a recessive manner and lead to the absence of enteric neurons, chromatophores, Schwann cells, sensory neurons and other trunk crest cell types [31,25,32]. Craniofacial features appear to be largely unaffected in zebrafish *sox10* mutants, which is thought to be due to compensation by the *SoxE* family member *sox9b* in ectomesenchymal neural crest [33,34]. *Sox10* has also been shown to play a continued role in the maintenance and differentiation of adult melanocyte stem cells in mouse [35,36].

Melanoma, a highly aggressive form of cancer originating from neural crest-derived melanocytes, shows signs of melanocytes reverting to a crest like state as part of their disease progression [37–39]. Recently, *SOX10* has also been shown to play a crucial role in the overlapping identity of neural crest stem cells and melanoma, where silencing of *SOX10* suppresses the neural crest stem cell-like properties in melanoma [38]. Together these data indicate a role for the neural crest differentiation pathway in melanomagenesis.

Many crucial transcription factors involved in the neural crest GRN have been identified and studied in depth across a number of different species, but a lot of their downstream targets and interaction partners still remain to be elucidated. For example, TFAP2A ChIP-Seq analysis using human neural crest cells has identified over 4,000 potential TFAP2A binding sites and established TFAP2A as a chromatin initiating factor [40]. This large number of putative TFAP2A downstream targets now requires functional validation.

To identify novel effectors and temporal trajectories of the *tfap2a;tfap2c* and *sox10* neural crest network we used zebrafish mutants in genes required at different levels of neural crest differentiation in a single whole embryo transcriptomic screen across different developmental stages. This screen involved genotyping individual embryos at all relevant developmental stages before sequencing their mRNA. Using a high number of replicates has proven vital in identifying true neural crest signal while also highlighting genetic background effects such as haplotype-specific expression. Stepwise genetic ablation of *tfap2* signalling uncovers dose-dependent genetic compensation between paralogues. To validate novel candidates emerging from this analysis we applied a reverse genetics approach to knock out genes of interest using both ENU and CRISPR/Cas9 mutagenesis [41–43]. Taken together, this work has identified early activation of the neural crest GRN and the core gene set underlying genetic compensation of *tfap2a* or *tfap2c* perturbations. Our screen has also identified novel downstream neural crest genes and a role for maternal expression of the Hippo signalling member *yap1* in the differentiation of melanocytes. All resources are publically available and we envisage that this will lead to a deeper understanding of neural crest biology.

## Results

Our collection of mutations in previously well studied zebrafish mutants (*tfap2a*, *tfap2c*, *sox10* and *mitfa*) as well as a newly associated neural crest mutant (*yap1*–this study) encompasses an early undifferentiated, premigratory neural crest state through to terminal differentiation of different crest cell types, in particular the melanocytes.

### Neural crest GRN is initiated at genome activation

Neural crest cells can be readily identified as the first somites begin to form, however it is not clear when the neural crest GRN becomes active in the zebrafish embryo. We used a wild-type developmental time course we had published previously [44] encompassing 18 stages from zygote to 5 dpf to identify the specific time points at which relevant transcripts are activated and their expression over time. In addition, the use of single embryos reveals the natural variation across individuals (Fig. 1b-g). In zebrafish, the genome first becomes transcriptionally active between the 1K-Cell and Dome stage [45–47]. A number of early neural crest transcription factors - *foxd3*, *tfap2a*, *tfap2c* - can be detected at the Dome stage, which is much earlier than neural crest has typically been thought to be induced (Fig. 1b-d) [2]. Their downstream targets *sox9b* and *sox10* begin to be expressed between 75% epiboly and when the first somites appear (Fig. 1e-f). Both *sox9b* and *sox10* have been shown to be robust markers for premigratory neural crest cells in zebrafish [48].

### Identification of a neural crest–enriched gene set

We first created a catalogue of genes enriched in premigratory and differentiating neural crest cells as a reference set for the subsequent transcriptional analysis of the neural crest mutants. We used Fluorescence-Activated Cell Sorting (FACS) on dissociated cells from whole embryos of the *sox10:mg* line [49] at 22-23 hours post fertilisation (hpf). The transgenic reporter labels neural crest nuclei (mCherry) and crest cell membranes (GFP). At 22-23 hpf neural crest cells migrate along the anterior-posterior axis and their differentiation is more advanced at the rostral than caudal part of the embryo. We therefore reasoned that this stage would provide us with a comprehensive mixture of neural crest differentiation states. We observed a delay in the membrane bound GFP signal causing two separate neural crest populations; one labelled only with the nuclear mCherry marker, and a second labelled both with mCherry and the membrane bound GFP (Fig. 1h). We sorted these two populations separately along with a third non-transgenic population for pairwise differential expression analysis, however for the purposes of this study we pooled the neural crest cell data. Our second aim was to assess whether we could gain more information by investigating the transcriptional profiles of cranial crest and trunk crest separately. We separated heads and tails of embryos from the same stage and isolated a single neural crest population from each tissue type comprising both mCherry+ and mCherry+/GFP+ cells as well as an unlabelled non-crest population. All cell populations were processed to produce polyA RNA-Seq libraries and sequenced. We compared transcripts detected in the transgenic neural crest cell populations to the non-crest cells using DESeq2 to produce neural crest-enriched gene sets (Fig. 1i). For comparison to our whole embryo data we pooled the resulting gene lists from the individual FACS experiments to produce a set of 4995 genes enriched in any FACS neural crest cell population (Table 1).

**Table 1.** Sequencing data and pairwise comparison gene lists.

| Figure Location | Data Source | FigShare DOI |
| --- | --- | --- |
| Figure 1i | FACS RNA-Seq gene lists | [10.6084/m9.figshare.6106082](https://figshare.com/doi/10.6084/m9.figshare.6106082) |
| Figure 1j | DeTCT gene lists (*sox10*, *mitfa*, *yap1*) | [10.6084/m9.figshare.6106091](https://figshare.com/doi/10.6084/m9.figshare.6106091) |
| Figure 2d | DeTCT *tfap2a*; *tfap2c* gene lists | [10.6084/m9.figshare.6106091](https://figshare.com/doi/10.6084/m9.figshare.6106091) |
| Figure 2d | UpSet gene subsets & ZFA Enrichment | [10.6084/m9.figshare.6170417](https://figshare.com/doi/10.6084/m9.figshare.6170417) |
| Supplemental Figure 2b | RNA-Seq *tfap2a*; *tfap2c* gene lists | [10.6084/m9.figshare.6106079](https://figshare.com/doi/10.6084/m9.figshare.6106079) |
| Figure 4c | UpSet gene subset lists | [10.6084/m9.figshare.6170474](https://figshare.com/doi/10.6084/m9.figshare.6170474) |
| Figure 5b-g | MCL cluster gene lists | [10.6084/m9.figshare.6170651](https://figshare.com/doi/10.6084/m9.figshare.6170651) |

### *sox10* knockouts effects on downstream targets appear transcriptionally at 19 hpf

In order to establish the initiation of the *sox10* GRN, a downstream target of *tfap2a;tfap2c,* we first created a transcriptional loss of function time course of *sox10* and its target, *mitfa*, by comparing gene expression of homozygous mutants and siblings. Zebrafish *sox10* mutant embryos form premigratory neural crest cells in the trunk but these cells fail to migrate and properly differentiate while cranial crest remain largely unaffected [25]. Mutants of the *sox10* downstream target *mitfa* have mostly correctly differentiated neural crest but specifically lack melanocytes of the body while showing mild differences in the numbers of the other two pigment cell types xanthophores and iridophores [50].

Figure 1j is an overview of all experiments carried out using DeTCT (differential expression transcript counting technique) 3’ tag sequencing [51]. Although *sox10* is appreciably expressed at the 1-4 somites stage (~10 hpf) (Fig. 1f) it is only at the 19 somite stage (19 hpf) where we detected a reduction in the abundance of one of its downstream targets, *mitfa,* in *sox10* knockout embryos (Fig. 1j). The majority of genes changing in the *sox10* 4 somite, 15 somites and 19 somites stages are localised on chromosome 3, the same chromosome as *sox10,* and are an example of haplotype-specific expression signals (Figure 1j, Table 1). We also found similar signals for the *mitfa* mutants at 4 somites with a very strong enrichment for chromosome 6 at 24 hpf (Table 1). Haplotype-specific signals will be discussed later.

We next analysed enrichments of terms from the Zebrafish Anatomy Ontology (ZFA) associated with differentially expressed genes and plotted all time points with significant enrichments (Supplemental Fig. 1). As expected, we found a strong and specific melanocyte signal in both mutants across all time points, with *sox10* mutants also showing a strong enrichment at 24 hpf for xanthophores and iridophores. By 36 hpf we also found an enrichment for the terms peripheral nervous system and nervous system which is consistent with an established role for *sox10* in peripheral nervous system development [25]. Previous data [25] and our developmental time course show that the expression of *sox10* begins early, following the establishment of the first neural crest cells at about 4 somites. It is only at the 19 somite stage, however, in which we detect the first molecular signal via the reduction of *mitfa* transcript, and only at 24 hpf do we see the first ZFA enrichments.

### Transcriptomic profiling of neural crest genetic ablation at three developmental stages using 3’ tag sequencing

Based on the wild-type expression of *tfap2a* and *tfap2c*, the morphological double mutant phenotype and the *sox10* molecular phenotype we chose three time points, 4 somite, 15 somites and 24 hpf, for the transcriptomic screen of *tfap2a;tfap2c* mutants. At the 4 somite stage pluripotent neural crest stem cells should be well established based on *snail1b* expression [34] and detectable with a whole embryo transcriptomic approach.

To genetically ablate the neural crest, we created double carrier fish for *tfap2a^+/sa24445^;tfap2c^+/sa18857^* (denoted as *tfap2a^+/-^;tfap2c^+/-^* from here on) alleles, using mutants produced by the Zebrafish Mutation Project (ZMP http://www.sanger.ac.uk/resources/zebrafish/zmp/) [41]. We confirmed the phenotypes previously described in *tfap2a;tfap2c* depletion experiments [15,18]. Double homozygous embryos were indistinguishable from wild-type siblings at the 4 somites stage but were slightly elongated/dorsalised by the 15 somites stage and were clearly discernible by 24 hpf (Fig. 2a-b). Notably, we also identified a specific pattern of reduction of dorsal tail melanocytes in *tfap2a^−/-^;tfap2c^+/-^* embryos at 48 hpf (Fig. 2c) in addition to the melanocyte reduction previously noted in *tfap2a^-/-^* embryos which demonstrates a dosage effect of *tfap2c* heterozygosity on *tfap2a* homozygous mutants. All other genotypic combinations were indistinguishable from their wild-type siblings at 48 hpf with *tfap2a^-/-^* carriers progressing to present craniofacial defects at 72 hpf as previously described [14].

**Figure 2.**
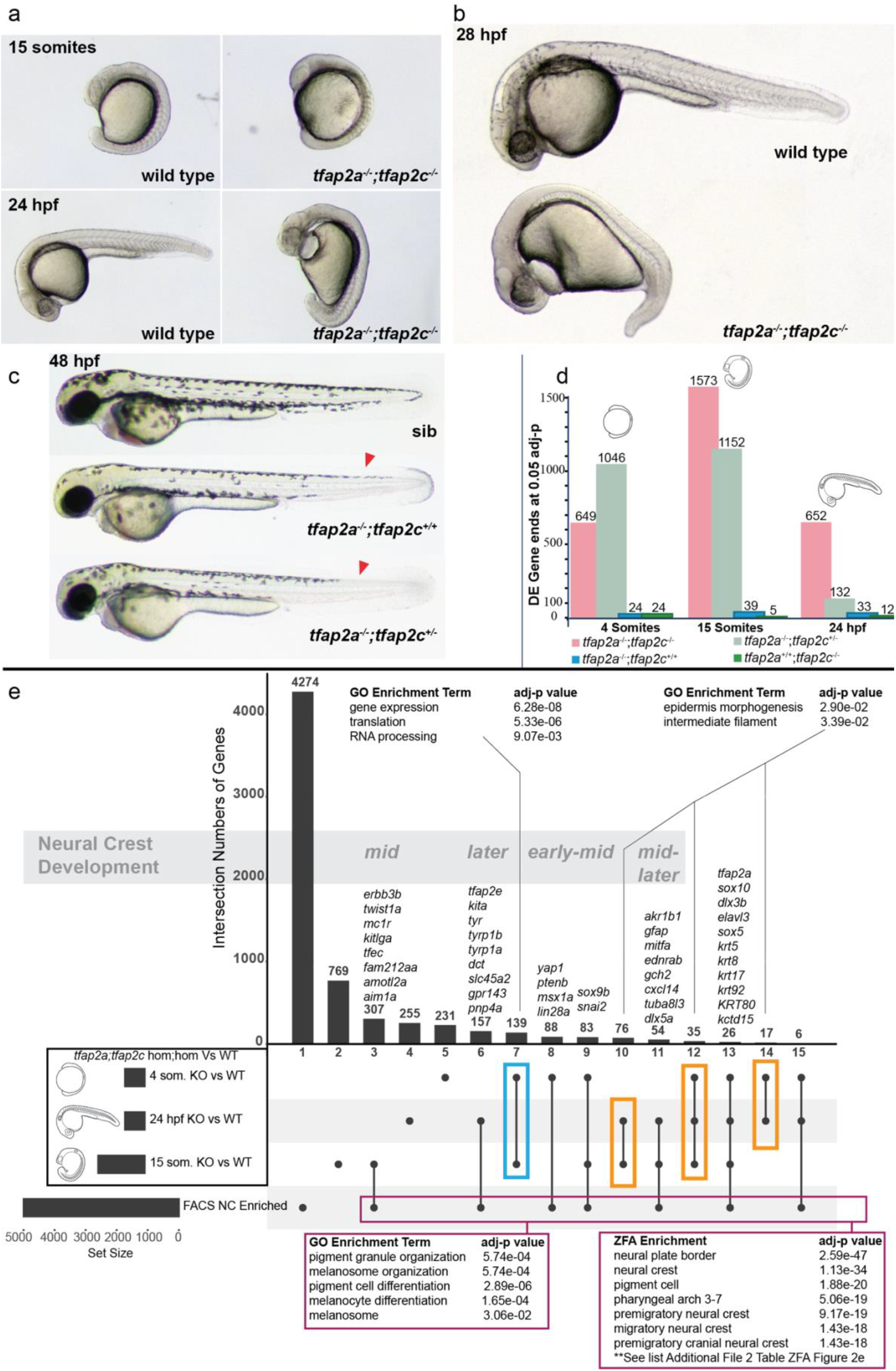
Molecular profiling of *tfap2a*;*tfap2c* mutants across multiple time points using 3’ tag sequencing. **a** *tfap2a^-/-^*;*tfap2c^-/-^* mutants present the first morphological phenotypes at the 15 somite stage. **b** By 28 hpf the morphological phenotype leads to an overall dorsalised form, bifurcation of the forming eye, heart oedema, and complete lack of neural crest cells. All other genotypes appear normal. **c** At 48 hpf the previously described reduction of melanocytes can be noted in *tfap2a^-/-^;tfap2c^+/+^* embryos and a modest reduction of melanocytes can be identified in the dorsal tail (red arrow heads) in *tfap2a^-/-^;tfap2c^+/-^* mutants. **d** Chart indicating the number of differentially expressed gene 3’ ends identified with an adjusted p-value of <=0.05 for each pairwise comparison of genotypes *tfap2a^-/-^;tfap2c^-/-^, tfap2a^-/-^;tfap2c^+/-^, tfap2a^-/-^;tfap2c^+/+^* and *tfap2a^+/+^;tfap2c^-/-^* to *tfap2a^+/+^;tfap2c^+/+^* siblings at 4 somites, 15 somites and 24 hpf **e** An UpSet diagram of the DE gene lists derived from the *tfap2a^-/-^;tfap2c^-/-^* vs. wild-type siblings (adj. p-value <0.05) for the 4 somites, 15 somite and 24 hpf stages and the list of neural crest enriched genes derived from sorted neural crest cells at 22-23 hpf. Individual subsets are marked with a black dot and overlaps with a connecting line. The total numbers of the subsets are represented with bars and number of genes above. GO enrichment was carried out on the subset found only in the 4 somites and 15 somites stages (blue box), the subsets indicated with the orange boxes and on all genes contained in the neural crest FACS enrichment and in at least one of the three different double knockout time points (magenta box). Using the developmental time course nature of the data allows for the grouping of the subsets into timing based on neural crest development starting with *early* neural crest specific gene expression and then moving towards *early-mid*, *mid*, *mid-later* and *later*.

In light of the observed phenotypes stemming from a dosage effect of *tfap2c* heterozygosity in *tfap2a* homozygous mutants our primary aim was to systematically investigate the genetic interactions of *tfap2a* and *tfap2c*. We therefore sequenced up to 10 embryos for all 9 genotypes at the three different stages to enable comparison of all genotypic combinations. We crossed double heterozygous *tfap2a;tfap2c* parents and collected embryos at the three developmental time points as single embryos. Following nucleic acid extraction and genotyping, single embryos were processed and global mRNA transcript levels determined using 3’ tag sequencing (Fig. 1j). After quality control and the removal of outlier samples we carried out pairwise analysis using DESeq2.

### Transcriptional phenotypes in *tfap2a* and *tfap2c* mutants differ greatly in magnitude when compared to their morphological outcomes

We first assessed how the transcriptomes of the different genotypic conditions behaved across time. Comparing the absolute numbers of differentially expressed (DE) genes of the four most relevant knockout genotypes over the three developmental time points revealed three major findings (Fig. 2d). Firstly, when compared to wild-type siblings, the number of genes differentially expressed in both *tfap2a* or *tfap2c* single homozygous embryos is very small in contrast to the double homozygous knockout and the *tfap2a^-/-^;tfap2c^+/-^* mutants indicating genetic compensation. Secondly, despite the severe morphological phenotype of double mutants at 24 hpf the number of DE genes was less than half of that at the 15 somites stage. Conversely, while only beginning to display a mild morphological phenotype at 48 hpf the *tfap2a^-/-^;tfap2c^+/-^* mutants showed a strong molecular phenotype at 4 and 15 somites, with a longer DE list at 4 somites than the double mutants. This molecular signature was strongly diminished by 24 hpf. Taken together this demonstrates that the complexity of transcriptional changes is not necessarily mirrored in the morphological phenotype, and vice versa.

### Overlapping multiple expression profiles groups genes by biological function

Next we analysed the transcriptional profile of complete ablation of the neural crest in *tfap2a^-/-^;tfap2c^-/-^* knockouts. A role for *tfap2a* has been previously described in both neural and non-neural ectoderm tissues which lead to the formation of the neural crest and the epidermis, respectively [15,52]. To separate transcripts into subsets specific to the neural crest or the epidermis we filtered the DE genes from the three developmental time points in *tfap2a^-/-^;tfap2c^-/-^* knockouts relative to wild-type siblings with the list of 4995 FACS-identified neural crest genes (Fig. 2e)[53]. When all genes which appear in at least one of the developmental stages and the neural crest FACS list are analysed together with their associated GO terms, there is an enrichment for pigment cells and melanocytes but no other neural crest subtypes (magenta box Fig. 2e). However, zebrafish anatomy enrichment (ZFA) returns a strong enrichment for the neural crest (Fig. 2e, Additional File 1). This finding highlights the current limitations of zebrafish GO annotation which has a bias for genes linked to pigmentation and lacks annotation for genes associated with earlier neural crest states.

A relatively small group of 26 genes appearing in all 4 data sets included *tfap2a*, *sox10* and many keratins. This could potentially signify an epidermal/neural crest precursor cell type which is in the process of committing to one of the lineages.

Comparison of the three developmental times points places genes into “early,” “mid,” and “later” neural crest-specific groups. Each of these groups contain numerous examples of previously characterised neural crest-specific genes which helps to validate this approach, but also many unannotated genes or genes previously not associated with the neural crest (Table 1).

The gene lists shared between the different stages but not found in the neural crest FACS data set (orange boxes Fig. 2e) and their Gene Ontology (GO) term annotation revealed an enrichment for epidermal-related terms. Another subset from the 4 somite and 15 somite stages that is not present in the NC-enriched gene list is a group of genes enriched for *expression*, *translation* and *RNA processing* (blue box Fig. 2e).

### *tfap2a;tfap2c* genetic compensation

Our next question was how the transcript levels of *tfap2a* and *tfap2c* along with three well characterised neural crest-specific genes (*foxd3*, *sox10* and *sox9b*) behaved across all 9 genotypes and the three developmental stages (Fig. 3a-o). At 4 somites, embryos homozygous for either *tfap2a* or *tfap2c* had significantly lower transcript abundances for their respective genes indicating that nonsense-mediated decay had most likely occurred (Fig. 3a-b)[54]. A genetic interaction is evident in *tfap2a^-/-^;tfap2c^+/+^* embryos between *tfap2a* and *tfap2c* with higher levels of wild-type *tfap2c* transcripts than in wild-type siblings (Fig. 3b) while *tfap2a* is not increased in the inverse case of *tfap2a^+/+^*;*tfap2c^-/-^* mutants (Fig. 3a). This indicates that already by the 4 somite stage, the neural crest GRN is able to detect reduced levels of *tfap2a* in knockouts and compensation by *tfap2c* is established. *foxd3* is significantly reduced in both *tfap2a^-/-^;tfap2c^-/-^* and *tfap2a^-/-^;tfap2c^+/-^* embryos compared to wild-type siblings, but not in *tfap2a^-/-^;tfap2c^+/+^* embryos, providing further evidence for a compensatory role of *tfap2c* when levels of *tfap2a* are reduced. Both *sox10* and *sox9b* behave in a similar manner to *foxd3* at 4 somites. These data also demonstrate that only one of the possible four *tfap2a* or *tfap2c* alleles is required to ensure early neural crest cell identity and differentiation.

**Figure 3.**
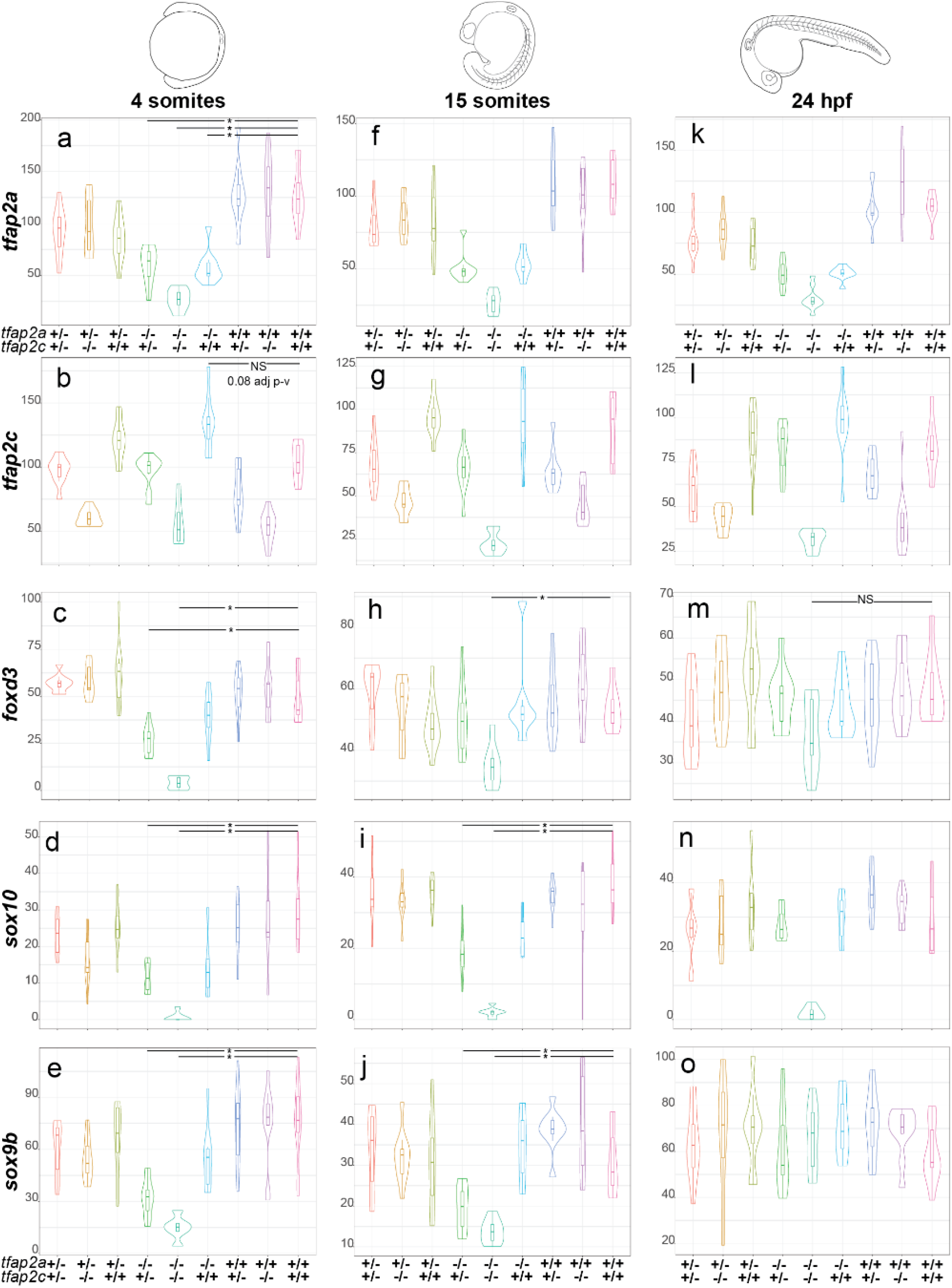
Expression of sox10, sox9b and foxd3 in *tfap2a;tfap2c* mutants across 3 developmental time points. Normalised counts and gene name to the left of the violin plots and the corresponding genotypes for *tfap2a* and *tfap2c* at the bottom. All plots are ordered by the time points shown on the top of the figure. **a** At 4 somites levels of *tfap2a* are significantly lower than in wild-type siblings in all *tfap2a^-/-^* genotypes. **b** Levels of *tfap2c* present at elevated levels in *tfap2a^-/-^*;*tfap2c^+/+^* embryos when compared to wild-type siblings but fail the statistical cut off with 0.08 adj p-value. **c-e** Levels of *foxd3*, *sox10* and *sox9b* are significantly different in both *tfap2a^-/-^*;*tfap2c^+/-^* and *tfap2a^-/-^*;*tfap2c^-/-^* embryos but not in *tfap2a^-/-^*;*tfap2c^+/+^*. **f-g** At 15 somites the levels of *tfap2a* and *tfap2c* recapitulate trends observed at 4 somites stage. **h** Levels of *foxd3* are only significantly different in *tfap2a^-/-^*;*tfap2c^-/-^* embryos when compared to wild-type siblings. **i-j** The levels of *sox10* and *sox9b* are both significantly different in *tfap2a^-/-^*;*tfap2c^+/-^* and *tfap2a^-/-^*;*tfap2c^-/-^* embryos compared to wild-type siblings. **k-l** The profiles of *tfap2a* and *tfap2c* at 24 hpf again remain similar to the two previous time points across all genetic combinations. **m** At 24 hpf the levels of *foxd3* are not significantly different across any genotypes. **n** The levels of *sox10* are markedly down in only the *tfap2a^-/-^*;*tfap2c^-/-^* embryos and levels of *sox9b* are unchanged across all genotypes **o**. Statistical significance of below 0.05 adj p-value is denoted with a *. Not all significant differences have been labelled. NS is to emphasise pairwise comparisons which fail an adj. p-value <0.05 cut off.

At 15 somites the transcriptional profiles of *tfap2a* and *tfap2c* remain similar to the 4 somite stage (Fig. 3f-g). Levels of *foxd3, sox10* and *sox9b* all remain significantly reduced in *tfap2a^-/-^ tfap2c^-/-^* embryos (Fig. 3h-j) while in *tfap2a^-/-^tfap2c^+/-^* embryos *foxd3* levels have begun to recover but expression of *sox10* and *sox9b* is still reduced.

By 24 hpf the abundance of *tfap2a* and *tfap2c* remains much the same as the previous developmental stages (Fig. 3k-l). Interestingly, *foxd3* and *sox9b* levels are no longer significantly different in *tfap2a^-/-^;tfap2c^-/-^* embryos which is suggestive of their exit from the neural crest GRN or initiation of expression in non-neural crest tissues, but levels of *sox10* remain strongly reduced in the double mutants (Fig. 3m-o). Also, *tfap2a^-/-^;tfap2c ^+/-^* embryos now have levels of *foxd3*, *sox9b* and *sox10* comparable to wild type which suggests a general recovery of the neural crest GRN by this stage. These data show that the time point of the strongest molecular phenotype and *tfap2c* compensation is at around 4-15 somites with the morphological phenotypes beginning to emerge by 15 somites.

### RNA-Seq on *tfap2a;tfap2c* knockouts at 15 somites confirms 3’ tag sequencing data and produces a more detailed transcriptional landscape

To further investigate the dose-dependent compensation while also creating a more detailed transcriptomic profile of pluripotent and differentiating neural crest cells, we carried out RNA-Seq on *tfap2a;tfap2c* knockouts at the 15 somite stage. All 9 genotypes were assessed using a total of 90 single embryos. Principal component analysis highlights that *tfap2a^-/-^;tfap2c^-/-^* and *tfap2a^-/-^;tfap2c^+/-^* are most similar on a molecular level in spite of their vastly different morphological phenotypes (Supplemental Fig. 2a). Pairwise comparisons of four different genotypes to their wild-type siblings shows high numbers of genes changing in both *tfap2a^-/-^;tfap2c^-/-^* and *tfap2a^-/-^;tfap2c^+/-^* groups (Supplemental Fig. 2b, Table 1). The majority of significant genes have reduced transcript levels in double mutants with robust p-values (Supplemental Fig. 2c). The 15 somite 3’ tag sequencing and RNA-Seq data sets showed good correlation of the detected DE genes at an adjusted p-value < 0.01 (Supplemental Fig. 2d)

Hierarchical clustering on the significantly changed genes from the *tfap2a^-/-^;tfap2c^-/-^* versus wild type pairwise comparison and ZFA enrichment placed genes into functional groups. While loss of both *tfap2a* and *tfap2c* leads to a reduction in genes involved in neural crest and epidermis development it also leads to an upregulation of genes associated with neural terms (Supplemental Fig. 2e).

### Identifying genes required for *tfap2a;tfap2c* knockout compensation

The 3’ tag sequencing analysis had highlighted that both *tfap2a^-/-^;tfap2c^-/-^*and *tfap2a^-/-^;tfap2c^+/-^* gave the most extensive molecular phenotypes, but *tfap2a^-/-^;tfap2c^+/-^* were morphologically indistinguishable from wild-type siblings at 15 somites whereas *tfap2a^-/-^;tfap2c^-/-^* presented obvious morphological phenotypes by that stage. Hence a single allele of *tfap2c* is sufficient to rescue the morphological *tfap2a^-/-^;tfap2c^-/-^* neural crest specification and differentiation phenotype despite the observed effect on the transcriptional level. We were therefore keen to understand which genes are involved and may be required for the rescue of the morphological phenotype.

First, we assessed expression of *tfap2c* in the RNA-seq data and found that the levels of *tfap2c* were significantly higher in *tfap2a*^-/-^ embryos at 15 somites when compared to wild-type embryos (Fig. 4a). We then compared the sets of differentially expressed genes derived from the pairwise comparisons of wild type with *tfap2a^-/-^;tfap2c^-/-^* and *tfap2a^-/-^;tfap2c^+/-^*, respectively. The vast majority of DE genes in the *tfap2a^-/-^;tfap2c^+/-^* condition were also changed in the *tfap2a^-/-^;tfap2c^-/-^* embryos (Fig. 4b). If we consider that there is a total of four alleles between *tfap2a/c,* this demonstrates that loss of a third *tfap2a/c* allele affects the neural crest GRN, however the transcriptional changes are not sufficient to derail neural crest specification and differentiation. Crucially, this identifies a core set of *tfap2a/tfap2c* responding genes, separate from secondary downstream events caused by differentiation failure and tissue loss.

**Figure 4.**
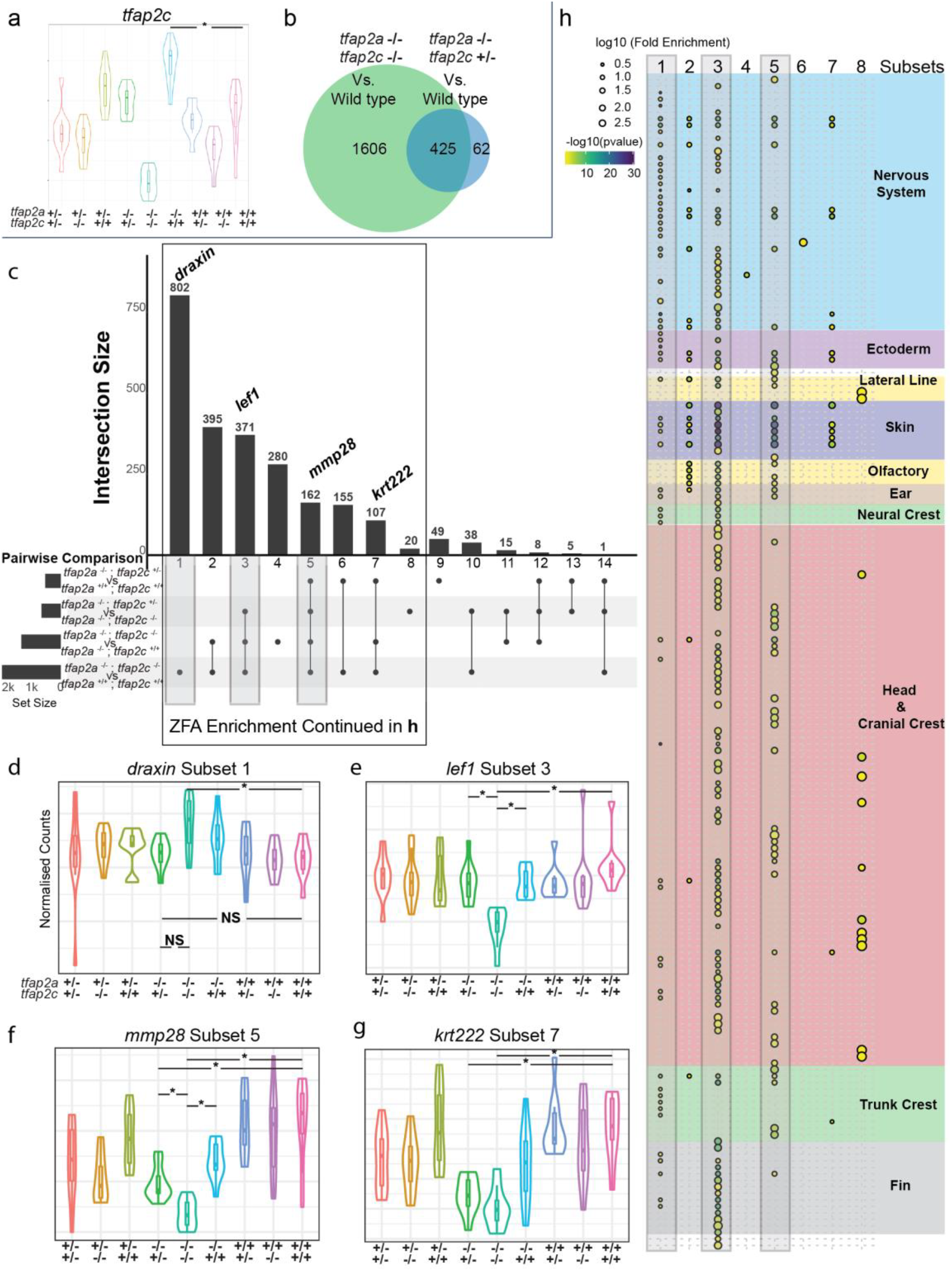
Identification of NC specific gene subsets in *tfap2a;tfap2c* mutant RNA-Seq 15 somite data. **a** RNA-Seq at 15 somites, an * indicates a significant (adj. p-value <0.05) increase of *tfap2c* transcript in *tfap2a^-/-^*;*tfap2c^+/+^* embryos when compared to wild-type siblings. **b** Overlapping gene lists comparison of significantly (adj. p-value <0.05) differentially expressed genes when *tfap2a^-/-^*;*tfap2c^-/-^* and *tfap2a^-/-^*;*tfap2c^+/^*^-^ are compared to wild type siblings. **c** Subsetting of gene lists from four different pairwise comparisons. The subsets are labelled 1-14 and the genes from **d-g** labelled at the top of the groups they belong to. Groups 1, 3 and 5 have grey boxes around their overlapping groups. **d-g** Examples of violin plots for the four subset groups with “*” signifying a <0.05 adj p-value between two groups and NS indicating not significant. Genotypes of the embryo groups are listed at the bottom of each plot. **g** Subsetting of gene lists from four different pairwise comparisons. The subsets are labelled 1-14 and the genes from **c-f** labelled at the top of the groups they belong to. Groups 1, 3 and 5 have grey boxes around their overlapping groups. **h** ZFA enrichment was carried out on all 14 subsets but only returned significant enrichment for groups 1-8. The log_10_[Fold Enrichment] is designated by the size of the circle and the colour represents-log_10_[p-value]. Grey bars correspond to the same subsets in **g**. Anatomy terms have been manually organised based on the themes to the right. The actual terms have been cropped and placed in (Supplemental figure 3 ZFA Enrichment) for ease of reading.

Next we dissected the full ablation response (*tfap2a^-/-^;tfap2c^-/-^*) using the partial ablation profiles (*tfap2a^-/-^;tfap2c^+/+^ and tfap2a^-/-^;tfap2c^+/-^*). As a single allele of *tfap2c* is able to maintain neural crest specification we sought to identify genes that are sensitive to different levels of *tfap2c*. To this end we ran four differential gene expression (DGE) analyses: double homozygous embryos against embryos with one or two wild-type alleles of *tfap2c*, and wild-type embryos against *tfap2a^-/-^;tfap2c^-/-^* or *tfap2a^-/-^;tfap2c^+/-^*. Next we overlapped the four lists to produce 14 subsets (Fig. 4c). This identified several expression profile classes. Subset one contains genes where *tfap2a^-/-^;tfap2c^-/-^* knockout resulted in a mild, but significant change from wild-type siblings but there is no significant difference between *tfap2a^-/-^;tfapc^+/-^* and *tfap2a^-/-^;tfap2c^-/-^* or wild-type siblings, respectively, as is the case for *draxin* (Fig. 4d). For genes in subset three a complete *tfap2a^-/-^;tfap2c^-/-^* knockout resulted in a significant change from wild-type siblings however a single allele of *tfap2c* was sufficient to return the expression to wild-type levels. An example of this case would be *lef1* (Fig. 4e). Subset five contained genes that are only partially rescued. A single or even both wild-type alleles of *tfap2c* are unable to return expression to wild-type levels but the expression is still *significantly* different from the *tfap2a^-/-^;tfap2c^-/-^* condition, as exemplified by *mmp28* (Fig. 4f). Finally, subset 7 contained genes that are only rescued by two alleles of *tfap2c*, such as *krt222* (Fig. 4g).

### ZFA enrichment confirms specific neural crest signatures

We carried out ZFA enrichment on all 14 gene subset lists and obtained significant enrichments for subsets 1-8 (Fig. 4h, Supplemental Fig. 3). Subset three, the genes fully rescued by either one or two alleles of *tfap2c*, showed the strongest enrichment for terms associated with the neural crest, head and cranial crest and also fin. While fin enrichment may seem nonsensical for a 15 somite embryo this is due to the fact that many genes annotated for fin development are also involved in craniofacial development. A similar enrichment profile resulted from subset five, the genes where either one or two alleles of *tfap2c* rescued expression levels to a significant extent, but not completely. By contrast, the two largest subsets, containing genes that change in double homozygous embryos with respect to wild types, but not compared to *tfap2a^-/-^;tfapc^+/-^,* showed a bias towards nervous system and ectoderm enrichment. Crucially, subsets six and seven with genes that failed to be rescued by either one or two *tfap2c* alleles, had no or very little neural crest enrichment. This suggests these genes represent *tfap2a* targets outside of neural crest differentiation.

Taken together the enrichment analysis breaks down the full *tfap2a/tfap2c* knockout response into separate expression classes with different functional profiles. Subsets three and five contain genes that are fully or partially rescued by *tfap2c*, show the strongest neural crest enrichment and are thus most likely to represent the core of the *tfap2* neural crest GRN.

### Markov clustering reveals widespread haplotype-specific gene expression

Next we applied an expression correlation network and Markov clustering approach using Biolayout *Express^3D^* [55,56] to identify co-expression profiles independent from condition-driven differential expression analysis. We constructed a network graph with genes as nodes and their Pearson correlation as edges from the *tfap2a;tfap2c* RNA-Seq dataset and used Markov clustering (MCL) to divide the network into discrete sets of co-expressed genes. This identified *tfap2a* and *tfap2c*-specific components (Fig. 5a) within the larger co-expression network. The majority of the network clusters was dominated by co-expression groups of genes that share a genomic locus (Fig. 5a’ and Supplemental Fig 4). This is most likely driven by haplotype-specific expression which is extensively documented and studied in tissue- or cell-specific RNA-Seq data, but not in a whole organism context [57,58]. This suggests that the high genetic variability in zebrafish has a direct and widespread bearing on gene expression levels. In DGE analyses where homozygous mutants are compared to siblings haplotype-specific expression can lead to an enrichment of DE genes on the same chromosome as the mutation. Examples of such regions on chromosome 24, close to *tfap2a,* as well as on chromosomes 15 and 16, can be found in Supplemental Figure 4. We also identified the same phenomenon when analysing early time points in *sox10*, *mitfa* and *yap1* mutants (Figure 1j) where the majority of changing genes were on the same chromosome as the mutant being investigated. We have previously also noted the same effect across many different mutations and sequencing platforms.

**Figure 5.**
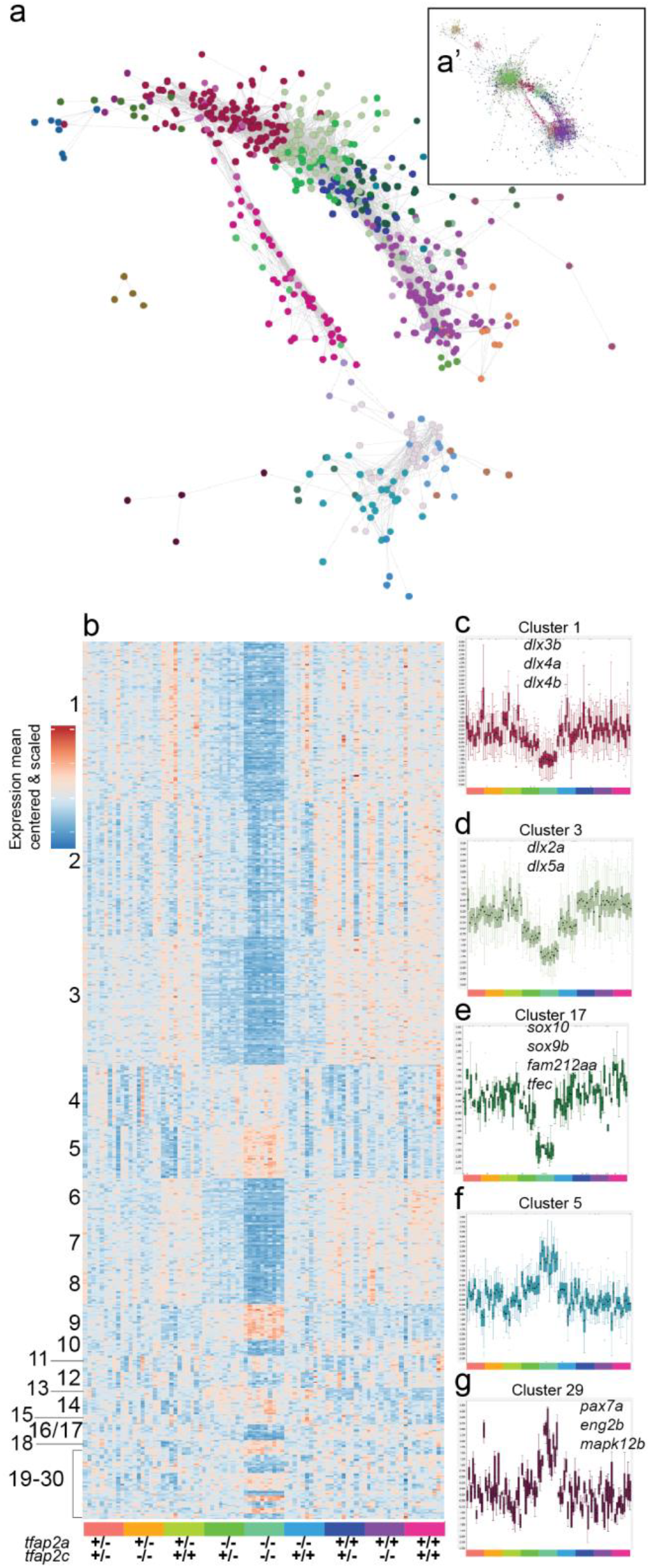
Network analysis and Markov clustering of RNA-Seq 15 somite data set. **a** Interaction network analysis of entire RNA-Seq 15 somite (0.7 Pearson correlation) data set represented as a subset. The entire interaction network can be found in **a’**. Each node represents a single gene and its colour corresponds to its cluster group. **b** A heatmap representing 24 MCL network clusters organised by cluster size and by genotype at the bottom. **c-g** Examples of individual clusters displayed as boxplots of the values for all the genes in the cluster (mean cantered and variance scaled). Samples are arranged as in **b** and are colour coded at the bottom of each cluster. Cluster number corresponds to the same cluster in **b**. Some genes contained in clusters are labelled on the plot.

### *tfap2a-* and *tfap2c-*specific gene clusters

Using MCL clustering we identified 30 clusters containing a total of 600 genes that were driven by changes in the *tfap2a* or *tfap2c* genotypes and organised the clusters into a mean centred and scaled heatmap (Fig. 5b). It is important to point out that in the previous analysis we compared lists derived from pairwise DGE comparisons, whereas MCL clusters all expressed genes based on their expression similarity across all samples. Therefore, these clusters might exclude genes that are identified in the DESeq2 analysis because of low expression correlation with other genes, but also include highly correlated genes which did not produce a significant adjusted p-value in the DESeq2 analysis.

The unsupervised clustering confirmed the strong signal in the double homozygous fish (clusters one and two) and dose-dependent compensation by *tfap2c* (cluster three). However, in addition it provided increased functional resolution. For example, cluster 17 (Fig. 5e) was highly specific to neural crest effectors containing the *soxE* paralogues *sox10* and *sox9b*, the micropthalmia bHLH transcription factor *tfec* as well as the Pak4 kinase inhibitor *fam212aa* in addition to one uncharacterised gene (*si:ch211-243g18.2; ENSDARG00000044261*).

The differentiation of the neural crest also requires the down-regulation of specific groups of genes, for example to repress a neural fate. Cluster five (Fig. 5f) contains a collection of *soxB* family genes (*sox3*, *sox19a*, *sox19b*, *sox21b*), one of which (*Sox19)* being one of the earliest CNS markers in vertebrates [48]. Cluster five also includes another example of paralogues of *oct-*related transcription factors *pou3f2b* (*Oct-2*) and *pou3f3a*, which are associated with controlling CNS development. Cluster 29 (Fig. 5g) contains a collection of genes (*pax7a*, *eng2b*, *mapk12b* and *enfa2a*) which, based on the midbrain/hindbrain expression patterns of *pax7a* and *eng2b*, also suggests a developmental CNS role. All gene lists of individual clusters along with GO and ZFA enrichments can be found here (Table 1).

Using many replicates of single, genotyped, embryos from the same clutch has allowed us to identify haplotype-specific signals on a genome-wide scale. With a single allele of *tfap2c* sufficient to maintain a minimal neural crest GRN, we have compiled functional subsets of maintained genes, many of which are still poorly described and previously have never been associated with the neural crest. We have identified multiple cases where gene families or paralogues behave in the same manner, highlighting more potential examples of the compensatory nature of the GRN in general. To validate the association of novel genes with neural crest biology, we next analysed a set of candidates using a knockout approach.

### Validation of novel neural crest transcripts

We have identified a large number of novel neural crest candidate genes with poor or no functional annotation (Table 1). To validate a subset of these, we analysed the expression patterns or knockout phenotypes in zebrafish embryos. *wu:fc46h12; ENSDARG00000114516* transcripts were strongly reduced in a number of *sox10* mutant experiments (Table 1). To analyse where in the embryo *wu:fc46h12* is expressed, we performed *in situ* hybridisation on 24 hpf and 48 hpf wild-type and *sox10* mutant embryos. As a positive control for neural crest and xanthophores we included *in situ* hybridisation of *gch2*. At 24 hpf *wu:fc46h12* has an identical expression pattern to *gch2* in both wild-type and *sox10* mutants (Fig. 6a-d,g-h). At 48 hpf the expression of *wu:fc46h12* and *gch2* begins to diverge in wild types as *wu:fc46h12’s* expression domain becomes more specific to a ventral crest population (Fig. 6e-f), heart and dorsal aorta (Fig. 6e’-f’). The majority of these expression domains are also lost at 48 hpf in *sox10* mutants (Fig. 6g-h). We then created a CRISPR/Cas9 *wu:fc46h12^sa30572^,* but observed no obvious phenotype in homozygous embryos and raised homozygotes to adulthood. We carried out intercrosses of homozygous females with heterozygous males and observed heart oedema in maternal-zygotic (MZ) mutant *wu:fc46h12* embryos (Fig. 6i-j) but most larvae recovered and form swim bladders by 5 dpf.

**Figure 6.**
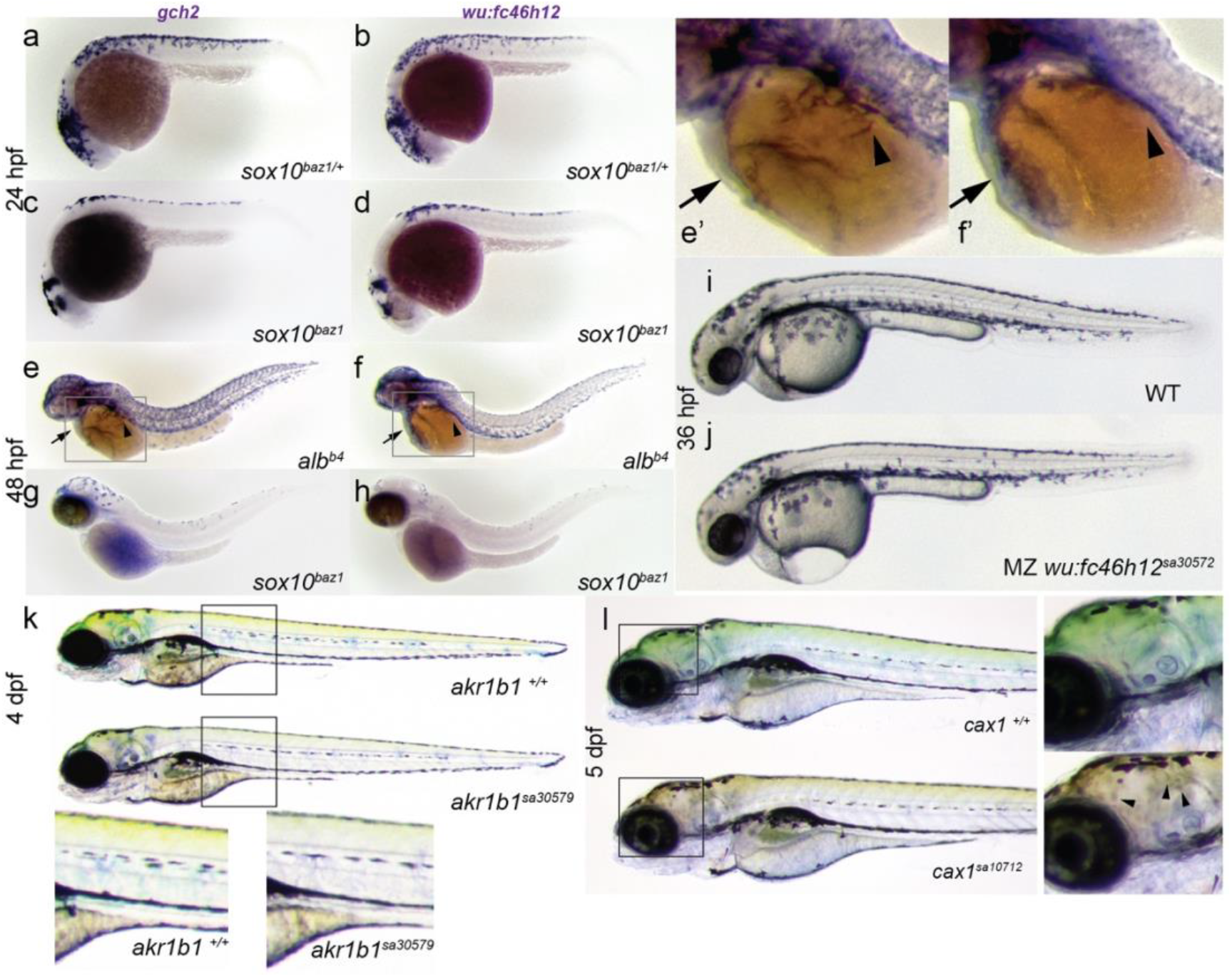
Functional analysis of pigment cell specific genes. **a-h** Whole mount *in situ* analysis of *wu:fc46h12* and *gch2* as a pigment cell comparison. *sox10^baz/+^* heterozygotes embryos as sibling controls **a-b** and mutant *sox10^baz1^* embryos at 24 hpf **c-d**. At 48 hpf *in situs* were carried out on *albino* embryos to serve as wild-type controls **e-f** with arrows indicating the heart and arrow heads the dorsal aorta. A blow up of this region can be found in **e’-f’**. **g-h** *E*xpression of *gch2* and *wu:fc46h12* at 48 hpf in *sox10^baz1^* mutants. **i-j** Wild-type and MZ*wu:fc46h12^sa30587^* embryos at 36 hpf with oedema around the forming heart **j**. **k** Wild-type sibling and mutant *akr1b1^sa30579^* at 4 dpf with mutant larvae presenting a reduction of yellow colour produced by xanthophores. Magnifications indicated with a black box. **l** Wild-type sibling and mutant *cax1^sa10712^* larvae at 5 dpf. Close ups indicated by black boxes around the head show dull yellow colour and abnormal cell morphology in mutants (arrowhead).

Two genes, *akr1b1* and *cax1* were both differentially expressed in the *tfap2a;tfap2c* and *sox10* data sets (Table 1). Using CRISPR/Cas9 we created a premature stop in *akr1b1^sa30579^*. Homozygous *akr1b1^sa30579^* fish develop normally but presented pale xanthophores (Fig. 6k). A premature stop in *cax1* was already available from the Zebrafish Mutation Project. The *cax1^sa10712^* allele presents a dulling in the colouring of xanthophores (Fig. 6l) as well as a rounding up of the cell morphology where typically xanthophores are highly dendritic. Homozygous *cax1^sa10712^* adults are viable and fertile, but MZ*cax1^sa10712^* embryos fail to develop normally beyond the start of somitogenesis pointing to a role of *cax1* during early embryonic development (Supplemental Fig. 5).

### A role for the Hippo signalling pathway in the neural crest

Expression of the transcriptional regulator *yap1* was reduced in double homozygous embryos in our 4 somite *tfap2a;tfap2c* 3’ tag sequencing data (Table 1) and *yap1* was also enriched in neural crest FACS-sorted cells (Figure 2e). In light of this, we assessed the DE gene lists in the *tfap2a;tfap2c* knockout versus wild-type sibling comparison from the RNA-Seq data set at 15 somites and found that three members of the Hippo signalling pathway *fat2*, *lats2* and *yap1*, had significant negative log_2_ fold-changes (Fig. 7a). These data suggested a role for Hippo signalling in neural crest cells.

**Figure 7.**
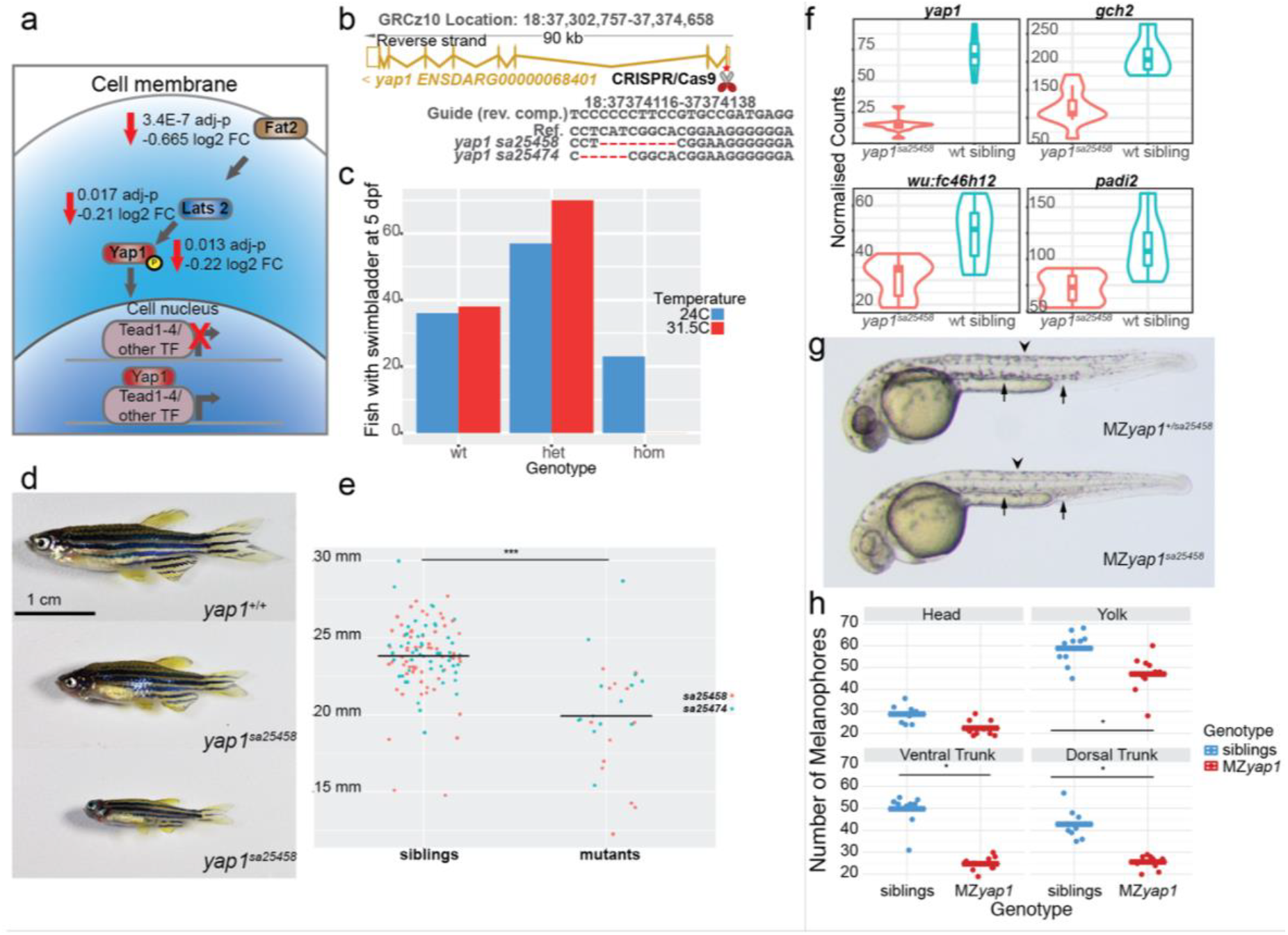
*yap1* mutants are temperature sensitive and play a role in melanocyte development. **a** Transcripts of members of the Hippo signalling pathway *fat2*, *lats2* and *yap1* were less abundant in *tfap2a;tfap2c* mutants when compared to wild-type siblings. A schematic showing their role in signal transduction and transcription inside of a cell. **b** Using CRISPR/Cas9 mutations were made in the first exon of *yap1* leading to the two alleles described. The exon-intron structure of the *yap1* transcript is shown in gold. The exact deletions are displayed below. **c** Embryos from a single clutch were split and raised at 24C and 31.5C with bars indicating the number of fish forming a swim bladder at 5 dpf grouped by *yap1^sa25458^* genotypes. **d** Homozygous *yap1* mutants are viable but present with a variation in size. **e** Quantification of size at two months of age with the corresponding genotypes for both *yap1* alleles. A statistically significant difference of <0.05 is indicated by “*”. **f** Normalised counts of 3’ tag sequencing data at 24 hpf comparing *yap1^sa25458^* mutants to wild-type siblings. All four genes, *yap1*, *gch2*, *wu:fc46h12* and *padi2* - have an adj. p-value <0.05. **g** Maternal zygotic *yap1* mutants present a strong reduction in melanocyte numbers at 36 hpf at both dorsal (arrow head) and ventral tail regions (arrow). **h** Quantification of melanocytes with the quantities on the left and then broken down into the regions of the head, yolk, ventral tail and dorsal tail. Each dot represents a region in a single larva, siblings in blue and MZ*yap1^sa25458^* in red. A statistical significance of <0.05 is indicated with “*”.

### *yap1* knockouts are temperature sensitive, homozygous viable and reduced in body size

To confirm a possible role for *yap1* in neural crest we targeted the first exon of *yap1* using CRISPR/Cas9 and created two alleles, *yap1^sa25458^* and *yap1^sa25474^*, leading to frame shifts and premature stops (Fig. 7b). When heterozygous carriers for either *yap1^sa25458^* or *yap1^sa25474^* were intercrossed and embryos raised at 28.5°C we were able to identify the previously described ocular phenotypes at 48-72 hpf in approximately 25% of embryos [59], albeit with variable penetrance depending on incubator temperature. Due to this observation, we tested whether these two *yap1* mutants were temperature sensitive. We split embryos from a single clutch and raised half at 24°C and the other half at 31.5°C. We then genotyped all fish which had formed a swim bladder by 5 dpf as larvae which fail to form a swim bladder by this time are not viable. Just over 20% of homozygous *yap1* mutant larvae formed a swim bladder when raised at 24°C but when raised at 31.5°C none of the homozygous *yap1* larvae formed a swim bladder (Fig. 7c).

To investigate whether fish raised at permissive temperature of 24°C were viable to adulthood, we raised intercrosses of *yap1* carriers for each allele (*yap1^sa25458^* & *yap1^sa25474^*) until 5 dpf then transferred them to our standard fish nursery. At 2 months post fertilisation, we observed that a subset of these fish were smaller than their siblings (Fig. 7d). We measured and genotyped intercrosses from both *yap1* alleles and confirmed that *yap1* homozygous fish were indeed smaller than their wild-type siblings (Fig. 7e).

### Zygotic *yap1* mutants show signs of neural crest GRN disruption

Although zygotic *yap1* mutants did not display obvious morphological phenotypes in neural crest cell types, we investigated whether there was a neural crest GRN phenotype by using a transcriptomic approach. We intercrossed *yap1^sa25458^* carriers, raised them at standard conditions of 28.5°C and collected embryos for 3’ tag sequencing at 4 somites, 15 somites and 24 hpf. The transcriptome profiles were normal at 4 somite and 15 somite stages, with the majority of the changed genes on the same chromosome as *yap1* (Figure 1j, Table 1). However, at 24 hpf the early xanthophore pigment cell marker *gch2* was significantly reduced in *yap1* mutants as well as *wu:fc46h12*, the newly identified pigment marker described above (Figure 7a-j). Interestingly, the early epidermis marker *padi2* was also reduced in *yap1* mutants (Fig. 7f).

### Loss of maternal *yap1* mRNA causes reduced melanocyte numbers at 30 hpf

Previous studies have shown a role for *yap1* in very early development of both zebrafish and medaka [60–62]. In zebrafish, this precedes zygotic genome activation and thus highlights a role for maternally deposited transcripts. The developmental time course data of *yap1* expression confirmed high levels of maternally deposited polyadenylated *yap1* (E-ERAD-475, www.ebi.ac.uk/gxa/home/).

Given the maternal deposition of *yap1* transcripts in the egg, we crossed heterozygous male *yap1^+/sa25458^* carriers to homozygous female *yap1^sa25458^* fish and evaluated the resulting MZ*yap1^sa25458^* larvae at the restrictive temperature of 31.5C. At approximately 30 hpf we observed a strong reduction in the number of melanocytes present in roughly half of the embryos. The previously described ocular phenotype [59] was also apparent in addition to a mild pericardial oedema (Fig. 5g). It is important to note that these larvae are otherwise morphologically stage matched. To confirm and quantify the melanocyte reduction we counted the number of melanocytes in four different sections; head, yolk, ventral trunk and dorsal trunk - of each larva and then genotyped them. A significant melanocyte reduction of about 50% in the yolk, ventral tail and dorsal tail was found with no major difference in the number of melanocytes in the head (Fig. 7h). This demonstrates that maternally deposited mRNA is able to rescue a melanocyte phenotype at 30 hpf further highlighting the very early induction of the neural crest GRN.

## Discussion

We have used transcriptional profiling on mutants affecting different steps of neural crest specification and differentiation to dissect the zebrafish neural crest GRN. We have used 3’ tag sequencing as a first pass screening method to then hone in with more detailed RNA-Seq. To make our data easily accessible to the research community we have placed the *tfap2a*;*tfap2c* 15 somite RNA-Seq experiment into Expression Atlas (www.ebi.ac.uk/gxa/homeexperimentE-MTAB-6106) for browsing and downloading and it will be made available with the next Expression Atlas release. The analysis of genotyped single embryos, independent from a visible phenotype, has allowed us to separate transcriptional responses from morphological outcomes. This approach is complementary to cell type-specific assays which require tissue manipulation and/or dissociation, much like the neural crest FACS RNA-Seq data set described here. Recently, elegant approaches have been developed to biotag specific cells *in vivo* and isolate their nuclei for further processing [63]. However, currently these methods require the pooling of embryos which would be challenging to apply to non-phenotypic embryos in loss of function analyses.

### Initiation of neural crest GRN before gastrulation, shortly following zygotic genome activation

The neural crest is typically described as being induced at the lateral edges of the neural plate after gastrulation. However, using wild-type developmental time course data we can place the activation of the neural crest transcription factors *tfap2a*, *tfap2c* and *foxd3* at the Dome stage, which follows zygotic genome activation and precedes gastrulation. In zebrafish, simultaneous loss of *tfap2a* and *foxd3* has been shown to genetically ablate the neural crest [64,65] with *tfap2a* and *foxd3* expressed in mutually exclusive compartments of the embryo at the shield stage, mid-way through gastrulation. The overlap of these expression domains forms the presumptive neural crest [65]. Recently in *Xenopus laevis* it has been shown that a high degree of overlap exists in the blastula pluripotent GRN and the neural crest GRN with the neural crest retaining the pluripotency of cells in the blastula stage rather than being induced later on in development [66]. Interestingly, the activation of the neural crest marker *crestin* also coincides with the Dome stage (E-ERAD-475, www.ebi.ac.uk/gxa/home/). This suggests that the establishment of the neural crest GRN, as it assumes its identity following blastula stages, shortly follows zygotic genome activation and places its initiation much earlier than previously shown in zebrafish and other vertebrates. This also raises the possibility of maternal mRNAs playing a larger role than previously thought in early neural crest initiation.

### Genetic ablation of the neural crest

In addition to *tfap2a;foxd3* loss of function a combined knockout of *tfap2a* and *tfap2c* genetically ablates the neural crest in zebrafish [15,65]. In the case of *tfap2a;foxd3*, *tfap2a* is thought to have an activator function whereas *foxd3* has been shown to act both as a repressor and an activator [67]. Knockouts of *tfap2a* fail to form normal jaws and have reduced numbers of melanocytes but still form neural crest cells. On a transcriptional level using 3’ tag sequencing, the number of genes which are differentially abundant in the *tfap2a* or *tfap2c* mutants alone are modest, 39 and 5 genes respectively at the 15 somite stage (Fig. 2d). At the 4 somite stage *tfap2c* acts in a compensatory manner as its overall abundance is increased by almost 50% in *tfap2a*^-/-^ embryos (Fig. 2b and Fig 5a). By removing a single *tfap2c* allele in *tfap2a^-/-^* embryos the number of changing genes jumps from 39 to 1152 (Fig. 2d), although this extensive change of gene expression is marked morphologically only with a mild decrease in the numbers of melanocytes in the tail at a much later stage. Using RNA-Seq at the 15 somite stage increases the total numbers of changing genes detected but the general trends remain much the same. *tfap2* family proteins are thought to form both homodimers as well as heterodimers [68]. This stepwise genetic ablation implies that *tfap2c* does not require *tfap2a* to initiate the early neural crest GRN and that either homodimers of *tfap2c* alone or potentially heterodimers with other *tfap2* family members are sufficient; however we do not see upregulation of any other *tfap2* genes.

### Dissection of the neural crest transcriptional network

*tfap2a* has been shown to play a role in the early stages of neural crest as well as the development of the epidermis. Both of these tissues arise at similar time points from ectoderm, and it is therefore crucial to separate the neural crest from the ectoderm signal. By combining multiple mutant data sets over developmental time along with the neural crest FACS data set we were able to establish the timing of when different levels of the neural crest GRN begin. Along with a large number of known downstream targets the subsets contain many uncharacterised genes, suggesting a role for these in pigmentation. We can further group genes which are more likely to not be specific to the neural crest but rather involved in epidermis development (Fig. 2e). Using the overlaps across the three different time points we have classified groups of genes from an “early” role to “mid” and then “later.” We have also further characterised trunk neural crest and melanocytes-specific downstream targets by analysing *sox10* and *mitfa* knockouts.

### Neural crest identity requires repression of a neural fate

The 15 somite stage had the highest number of differentially expressed genes in the *tfap2a*;*tfap2c* loss of function model and therefore we chose to investigate this stage in more detail using RNA-Seq. Using different subsetting approaches we have characterised distinct groups of neural crest genes and also have identified the core neural crest GRN that is maintained via *tfap2c*. The hierarchical clustered heatmap (Supplemental Figure 2e) highlights an enrichment of neural genes which are increased in the mutant samples. Considering the emerging model that neural crest cells are not actually induced *in situ* but rather a refinement of pluripotent blastula cells [66], our data support the notion that not only is the activation of the neural crest GRN important but also the repression of non-neural crest specific GRNs.

### Compensation of *tfap2a* knockout phenotypes via *tfap2c* and identification of genes involved in the neural crest rescue

RNA-Seq analysis of *tfap2a*;*tfap2c* knockouts and their siblings revealed an increase of *tfap2c* mRNA expression in *tfap2a* mutants at 15 somites. Although not addressed in this study, an interesting question now is: what is the molecular machinery which identifies the need for genetic compensation and how is it carried out? We find that whereas a single allele of *tfap2c* is able to rescue the early morphological neural crest ablation phenotype the expression of a core set of downstream effectors cannot be restored to wild-type levels. This separates the morphological phenotype, and its secondary molecular effects, from the primary gene-regulatory effect of *tfap2* loss of function. We have used this differential behaviour of downstream targets to identify genes which *tfap2c* is able to return to wild-type levels or to only partially rescue from the *tfap2a/c* double knockout. This confirmed known neural crest players but also added new genes to the neural crest GRN. The genes in subsets three and five (Fig. 4c-g) represent a core set of 371 and 162 genes, respectively, of the neural crest GRN required for early neural crest initiation and are most likely to be of high developmental and evolutionary importance.

### Genetic compensation via paralogues

Humans are particularly susceptible to haploinsufficient mutations in a number of neural crest-specific genes, including *sox10,* leading to Waardenburg syndrome or Hirschsprung disease, whereas the case in zebrafish seems to be different <u>[69]</u>. *sox10^+/-^* fish are adult viable and are phenotypically normal. Based on the developmental timing and clustering behaviour of the *soxE* family paralogues *sox10* and *sox9b*, there is a good probability that these two genes are able to compensate for each other in early neural crest cells. Similarly, fish with mutations in *tfap2c* are homozygous viable and *tfap2a^+/-^;tfap2c^-/-^* fish are indistinguishable from their wild-type siblings. By contrast, heterozygous mutations and alterations of *TFAP2A* lead to a number of developmental phenotypes in humans.

Previously, we and others have shown that the majority of mutations fail lead to an obvious morphological phenotype in the first 5 days of development in zebrafish [41,70]. Here, using the neural crest as a model we dissect the relationship between transcriptional robustness and morphological outcomes. Our study has also begun to reveal more evidence of genetic compensation in other paralogous genes. Unsupervised clustering has highlighted that entire gene families clustered together across development [44] and behaved in a similar manner in different genetic combinations in the *tfap2a*;*tfap2c* loss of function experiments (Figure 5b-e. Table 1).

Another example of possible paralogous compensation can be observed in the relatively mild developmental phenotypes of the *yap1* knockouts. Recently double knockouts of *yap1* and *taz (wwtr1)*, its paralogue, have shown much stronger early developmental phenotypes and are embryonic lethal [61]. A deeper understanding of genetic and functional paralogues with respect to mutual compensation versus division of function will provide mechanistic insight into gene function evolution.

### Identification of haplotype-specific signals

Use of high replicate genotyped samples has revealed an enrichment of differentially expressed genes on the chromosome carrying the mutation in the analysis of *sox10*, *mitfa* and *yap1* mutants (Figure 1j, Table1). This is most likely driven by stretches of homozygosity for the background linked to the mutation which produce different expression levels than the corresponding genomic loci in the control siblings. It is highly possible that some of these transcriptional differences could also have an effect on phenotypic outcomes of the mutation in question and could contribute to differences occasionally noted when a mutation is crossed into a different genetic background [71]. The differences in the haplotype-specific signals in the 4 somites and 24 hpf *mitfa* experiment also emphasise that additional factors such as stage and chromatin availability may be playing important roles. We can further demonstrate this effect on a genome-wide scale by increasing the total number of samples tested, as in the *tfap2a;tfap2c* 15 somite RNA-Seq experiment and allowing samples to cluster independently. In this analysis, we have identified groups of co-localised genes behaving in a similar manner on chromosomes other than the one carrying the mutation of interest. Although these clusters of genes are typically close to each other, the overall regions can span several hundred genes.

### A role for Hippo signalling in the neural crest

We have identified a reduction in the abundance of some Hippo signalling members in both our 3’ tag sequencing and RNA-Seq data sets. Previously, a role for Hippo signalling has been suggested in the neural crest using conditional mouse knockout models and in cell culture [72–74]. However, in the case of the mouse, complete *yap1* knockouts are not viable and in human iPS neural crest cell models both *YAP1* and *TAZ*(*WWTR*) require modulation. In zebrafish we show a role for maternally deposited *yap1* in the differentiation of melanocytes, however the effect on other neural crest subtypes remains to be investigated. Over the past few years post-embryonic neural crest stem cells have been identified in mouse and zebrafish [27,75,76]. The temperature sensitive *yap1* signalling model described here allows for the conditional inactivation of Hippo signalling and therefore the investigation of post-embryonic neural crest stem cells as well as other Hippo-dependent processes such as growth, pattern formation and regeneration later in development and in adults.

## Conclusions

Taken together, we have used transcriptional profiling and stepwise genetic ablation of the neural crest to divide the neural crest GRN into temporal and functional units containing new candidate genes alongside well known factors. The analysis of paralogue compensation separates the morphological neural crest ablation phenotype from the first expression changes to the core *tfap2* GRN. We confirm association of previously uncharacterised genes through knockout experiments and demonstrate a role of maternal transcripts in pigment cell development. Future studies of the functional gene clusters described here will help to further refine their role in neural crest development as well as their involvement in human genetic disorders and diseases such as neuroblastoma and melanoma.

## Materials and Methods

### Zebrafish Husbandry and Phenotyping of Mutants

Zebrafish were maintained at 23.5°C on a 14h light/10h dark cycle. Male and female zebrafish from genotyped heterozygous fish carrying mutations were separated overnight before letting them spawn naturally the next day. Fertilised eggs were grown at 28°C and single or multi-allelic phenotyping was carried out as previously described [41,77]. The *sox10^t3^* and *sox10^baz1^* alleles were a gift from Robert Kelsh and *mitfa^w2^* was previously a gift from Jim Lister [25,50].

### Embryo Collection

Embryos were either morphologically sorted into phenotypically abnormal and normal (*sox10^t3/baz1^*and collected at 28hpf, 36hpf and 48hpf) or collected blind at the stage of interest. Single embryos were placed individually into a well of a 2ml deep well block (Axygen, Cat number P-DW-20-C-S), snap frozen on dry ice and then stored at −80C.

### FACS

22-23 hpf embryos were collected from the zebrafish transgenic *sox10:mg* line which labels neural crest nuclei with mCherry and crest cell membranes with GFP. Dissociated cells were collected for FACS as previously described (Manoli et al., 2012). Briefly, embryos were dechorionated using 33 mg/ml pronase (Sigma) and pooled either as whole embryos or as pools of heads and tails. The yolks were removed using deyolking buffer (55 mM NaCl, 1.8 mM KCl, 1.25 mM NaHCO_3_) followed by digestion with trypsin-EDTA. Finally, the pellet was resuspended in FACSmax Cell Dissociation solution (AMS Biotechnology) and dissociated cells collected by passing the suspension through a 20 μm cell strainer (Sysmex Partec). Using appropriate gating, dissociated cells were sorted into mCherry positive, mCherry and GFP positive and unlabelled non-crest cells on the BD INFLUX. The data was analysed using FlowJo.

Sorted cells were collected and lysed in 110uls of RLT buffer (Qiagen) containing 1 μl of 14.3M beta mercaptoethanol (Sigma). The lysate was allowed to bind to 1.8 volumes of Agencourt RNAClean XP (Beckman Coulter) beads for 10 mins and RNA was eluted from the beads as per the manufacturer’s instructions. Total RNA was converted into cDNA libraries using the SMART-Seq V4 Ultra Low Input RNA kit (Clontech) followed by Nextera DNA Library Prep kit (Illumina) as per manufacturer’s instructions. Libraries were pooled and sequenced on Illumina HiSeq 2000 in 75 bp paired-end mode.

### Nucleic Acid Extraction

Frozen embryos were lysed in 100 μl RLT buffer (Qiagen) containing 1 μl of 14.3M beta mercaptoethanol (Sigma). The lysate was allowed to bind to 1.8 volumes of Agencourt RNAClean XP (Beckman Coulter) beads for 10 mins. The plate was then applied to a plate magnet (Invitrogen) till the solution cleared and the supernatant was removed without disturbing the beads. While still on the magnet the beads were washed thrice with 70% ethanol and RNA was eluted from the beads as per the manufacturer’s instructions. RNA was quantified using either Qubit RNA HS assay or Quant-iT RNA assay (Invitrogen).

### Genotype Confirmation

Genotyping was carried out according to [42]. Briefly, 1 μl of DNA from the extracted total nucleic acid was used to confirm the genotype of each sample using KASP SNP and InDel identification assays (LGC group) designed against our allele of interest. The genotyped plates were read on a plate reader (Pherastar, BMG Labtech) and 10-12 samples per genotype were selected for making libraries.

### Transcript counting

DeTCT libraries were generated as described previously [51]. Briefly, 300 ng of RNA from each genotyped sample were DNAse treated, fragmented and bound to streptavidin beads. The 3’ ends of the fragmented RNA were pulled down using a biotinylated polyT primer. An RNA oligo containing the partial Illumina adapter 2 was ligated to the 5’ end of the bound fragment. The RNA fragment was eluted and reverse transcribed using an anchored oligo dT reverse transcriptase primer containing one of the 96 unique index sequences and part of the Illumina adapter 1. The Illumina adapters were completed during a library amplification step and the libraries were quantified using either the BioPhotometer (Eppendorf) or Pherastar (BMG Labtech). This was followed by size selection for an insert size of 70-270 bases. Equal quantities of libraries for each experiment were pooled, quantified by qPCR and sequenced on either HiSeq2000 or HiSeq 2500.

Sequencing data were analysed as described previously [51]. Briefly, sequencing reads were processed with the DeTCT detag_fastq.pl script and aligned to the GRCz10 reference genome with BWA 0.5.10. The resulting BAM files were processed using the DeTCT pipeline, which results in a list of regions representing 3’ ends, together with a count for each sample. These counts were used for differential expression analysis using DESeq2 on pairwise combinations of samples. Each region was associated with Ensembl 86 gene annotation based on the nearest transcript in the appropriate orientation. False positive 3’ ends, representing, for example, polyA-rich regions of the genome, were filtered using the DeTCT filter_output.pl script with the --strict option, reducing the number of 3’ ends from 439,367 to 53943. Gene sets were analysed using topgo-wrapper for GO enrichment and Ontologizer for ZFA enrichment.

### RNA-Seq

Total nucleic acid was isolated from *tfap2a^+/sa24445^;tfap2c^+/sa18857^* intercrosses at 15 somites. KASP genotyping was used to identify all 9 possible genotypes. Total nucleic acid was treated with DNAseI (NEB, Catalogue number M0303L) and 10 replicates per genotype were processed. Ambion ERCC spike-in mix 2 (Cat. No. 4456740) was added to 200 ng RNA according to the manufacturer’s instructions and sequencing libraries were prepared using the Illumina TruSeq Stranded mRNA Sample Prep Kit. Libraries were pooled and sequenced on Illumina HiSeq 2500 in 75 bp paired-end mode.

Sequencing data were assessed using FastQC and aligned to the GRCz10 reference genome and Ensembl 86 transcriptome using TopHat2. Read counts per gene were generated using htseq-count and used as input for pairwise differential expression analysis using DESeq2. Gene sets were analysed using topgo-wrapper for GO enrichment and Ontologizer for ZFA enrichment. Custom R scripts were used for hierarchical clustering and principal component analysis. Count data were also clustered using Biolayout *Express^3D^*.

### Embryo and Fin Clip Genotyping

Genotyping of embryos and fin clips was performed as previously described [41,42]. Previously unpublished alleles used in this study are as follows:

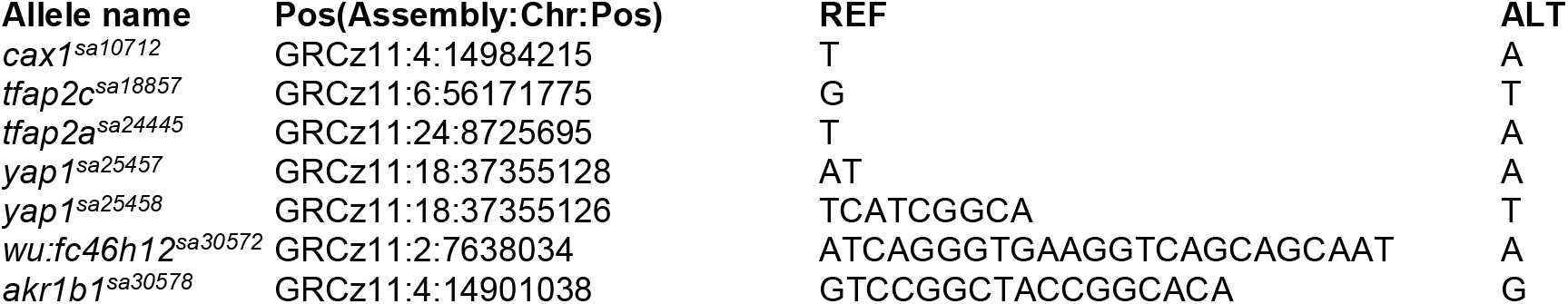

### RNA Whole Mount *In Situ* Hybridisation

RNA DIG-labelled probes were generated from cDNA libraries (Transcriptor High Fidelity cDNA Synthesis Kit, Roche) covering all relevant embryonic stages. PCR was performed and then TA cloned using TOPO-TA (Invitrogen). RNA riboprobes were produced using the T7- and SP6-promoter sequence, enabling synthesises of RNA in vitro transcription of the plasmid using T7- and SP6-RNA polymerase (Roche). All oligo nucleotide sequences are listed here:

wu:fc46h12_left1:CTGCTGACCTTCACCCTGATTCTG, wu:fc46h12_right1:GGTGTATTGCCTAAAACCCTCAGC wu:fc46h12_left2:ATTGCTGCTGACCTTCACCCTGAT, wu:fc46h12_right2:ATTGCCTAAAACCCTCAGCTTCCA.

### CRISPR/Cas9

Creation and identification of CRISPR/Cas9 zebrafish alleles were conducted as previously described using the zebrafish codon optimised double NLS Cas9 [78,43].

## Abbreviations

ALT: Alternative
Chr: Chromosome
dpf: days post fertilisation
DE: Differentially Expressed
DGE: Differential Gene Expression
DeTCT: Differential Transcript Counting Technique
ENA: European Nucleotide Archive
ENU: N-ethyl-N-nitrosourea
FACS: Fluorescence-Activated Cell Sorting
GFP: Green Fluorescent Protein
GRN: Gene Regulatory Network
hpf: hours post fertilisation
MCL: Markov Clustering
mut: mutant
MZ: Maternal Zygotic
NC: Neural Crest
NLS: Nuclear Localisation Sequence
PCA: Principal Component Analysis
PCR: Polymerase Chain Reaction
REF: Reference
sib: sibling
WT: Wild type
ZFA: Zebrafish Anatomy Ontology

## Acknowledgements

This work was supported by the Wellcome Trust [098051 and 206194]. The authors would like to thank Robert Kelsh for sharing the *sox10^t3^* and *sox10^baz1^* mutants and Claudia Linker for the *sox10:mg* transgenic line. The authors would also like to thank Rob Cornell and lab for discussions about the project along the way. We are grateful to all current and past Vertebrate Genetics and Genomics group members for advice and help, and Catherine Scahill and Nicole Staudt for manuscript feedback. The authors would also like to thank Nicola Goodwin and the RSF as well as sequencing pipelines at the Wellcome Sanger Institute for their excellent support and zebrafish care.

## Declarations

### Ethics Approval and consent to participate

Zebrafish were maintained in accordance with UK Home Office regulations, UK Animals (Scientific Procedures) Act 1986, under project licences 80/2192, 70/7606 and P597E5E82. All animal work was reviewed by The Wellcome Trust Sanger Institute Ethical Review Committee.

Consent to participate not applicable.

### Consent for publication

Not applicable

### Availability of data and material

The datasets supporting the conclusions of this article are available in ENA (https://www.ebi.ac.uk/ena). Accessions for all sequencing can be found in the Additional File 2 Data Sequence Archive Metadata.txt. DGE lists, clusters etc. described in this manuscript are deposited in a figshare collection (See Table 1 and 10.6084/m9.figshare.c.4077302). Zebrafish mutant lines will be made available upon request.

## Authors’ contributions

Conceptualization: CMD, EBN

Data curation: IMS, RJW, NW

Formal analysis: IMS, RJW, JEC, NW, CMD

Funding acquisition: DLS, EBN

Investigation: CMD, NW

Resources: DLS, EBN

Software: IMS, RJW

Supervision: EBN

Visualization: IMS, RJW, NW, CMD

Writing – original draft: CMD, EBN

Writing – review & editing: CMD, NW, RJW, IMS, JEC, EBN

---

**Supplementary Figure 1.**
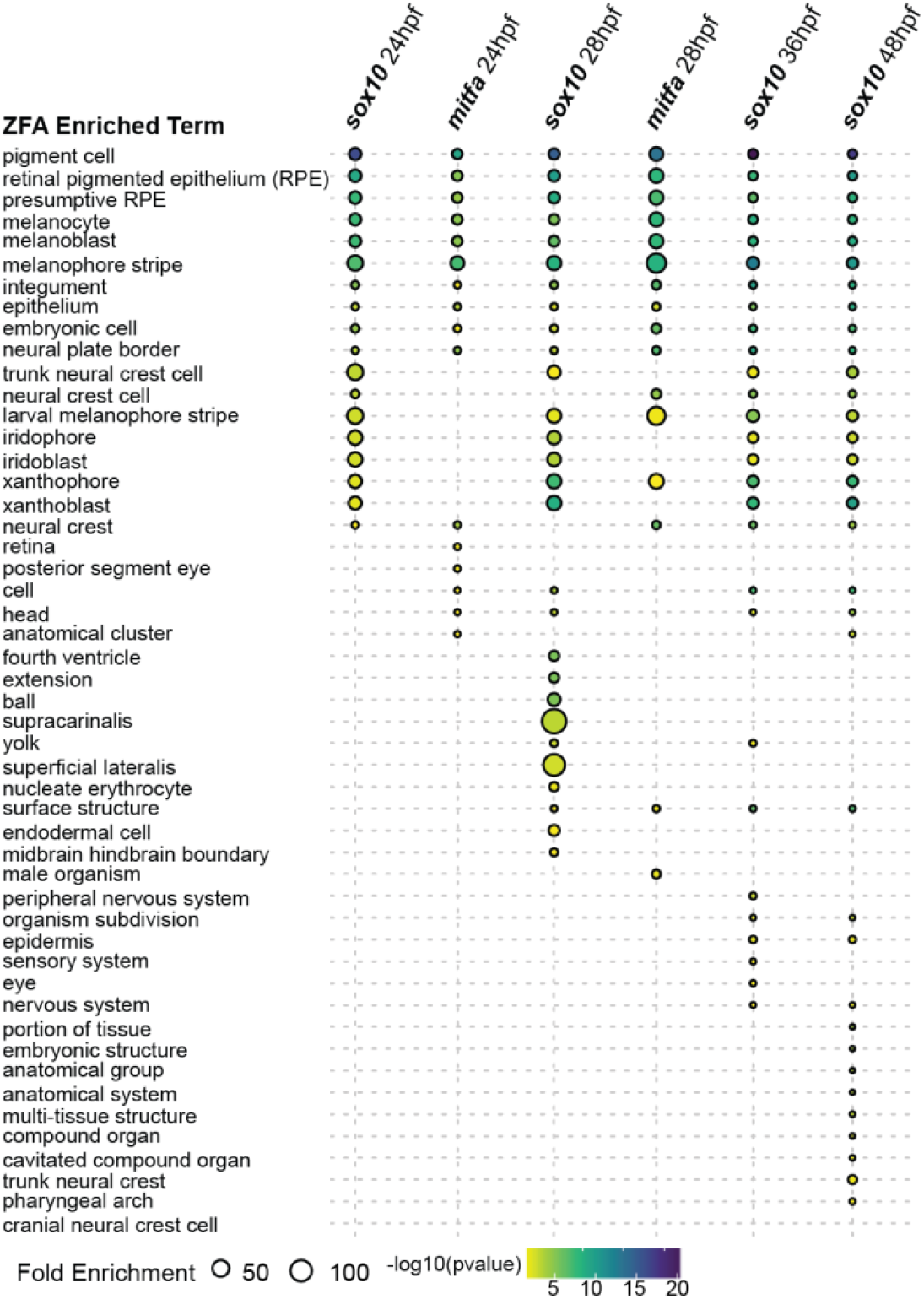
Zebrafish anatomy enrichment of *sox10* and *mitfa* mutants across multiple developmental time points ZFA enrichment was tested for all *sox10* and *mitfa* mutants compared to wild-type siblings at all time points shown in Figure 1j but only time points at 24 hpf or later returned significantly (adj. p-value <0.05) enriched terms.

**Supplementary Figure 2.**
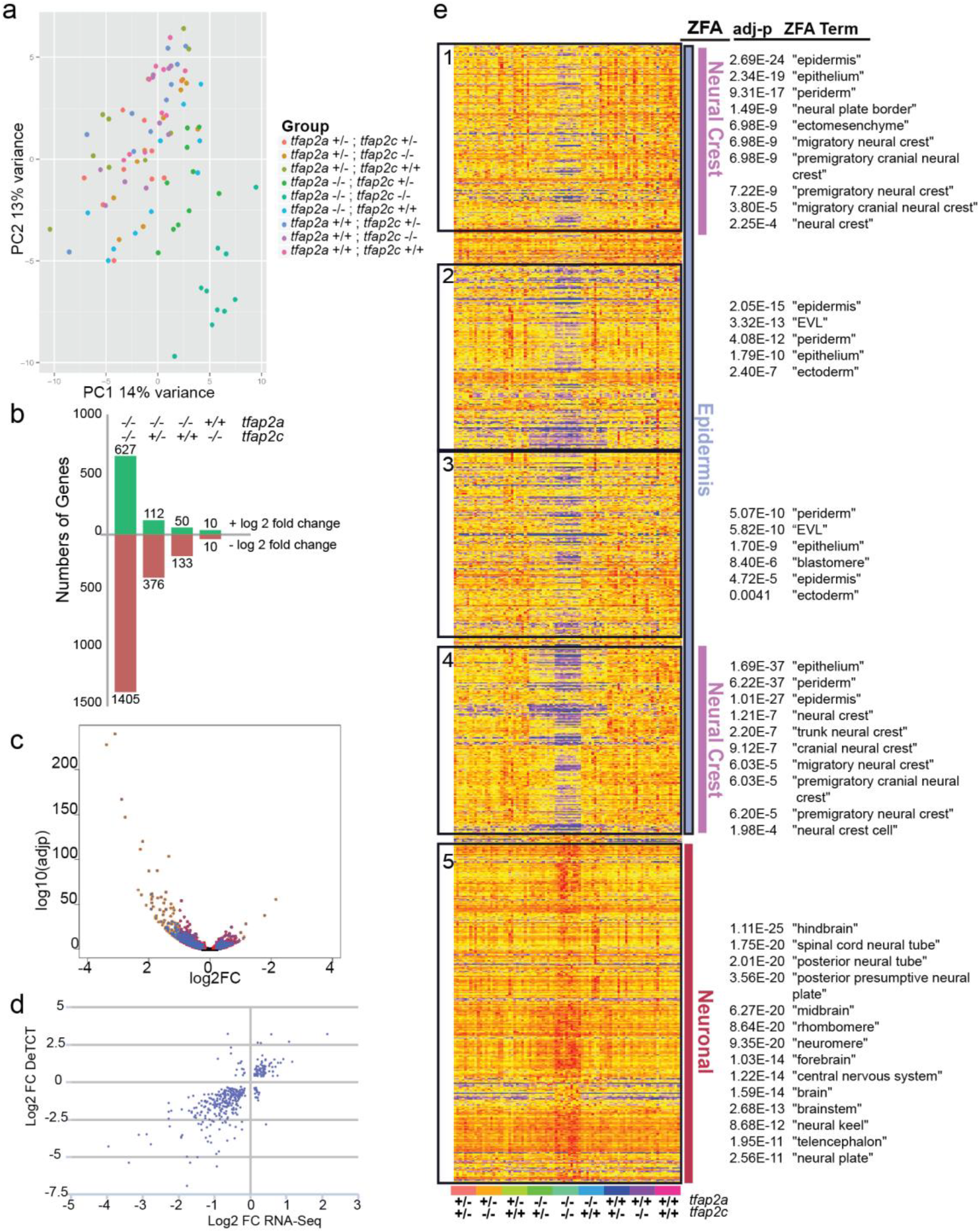
RNA-seq transcriptomic analysis of *tfap2a;tfap2c* mutants at 15 somite stage. **a** Principal component analysis of replicates of all 9 *tfap2a;tfap2c* genotypes showing the first two principal components. Dots representing a single embryo and genotype denoted by colour. **b** Bars denote the numbers of genes for four most relevant pairwise combinations (adj. p-value <0.05) with the numbers of genes with a positive log_2_ fold change in green and negative in red. The specific genotypes of *tfap2a* and *tfap2c* are listed across the top for each bar. **c** A pairwise comparison of RNA-seq of *tfap2a^-/-^*;*tfap2c^-/-^* versus their wild-type siblings at 15 somites. The adj p-value on the y axis and the log_2_ fold change on the x axis. **d** Comparison of 3’ tag sequencing (y axis) and RNA-Seq (x axis) log_2_ fold change of genes with an adj p-value <0.01 in the *tfap2a^-/-^*;*tfap2c^-/-^* versus wild-type siblings pairwise comparison showing an overall linear correlation. **e** Heatmap of gene expression with an adj p-value <0.05 from *tfap2a^-/-^*;*tfap2c^-/-^* to wild-type siblings pairwise comparison. Genes are hierarchically clustered with the samples organised by genotype. The heatmap was broken into five blocks shown in black boxes and ZFA enrichment was carried out on those blocks. ZFA enrichments with their corresponding significances are depicted on the right. ZFA terms were further broadly categorised into epidermis, neural crest and neuronal.

**Supplementary Figure 3.**
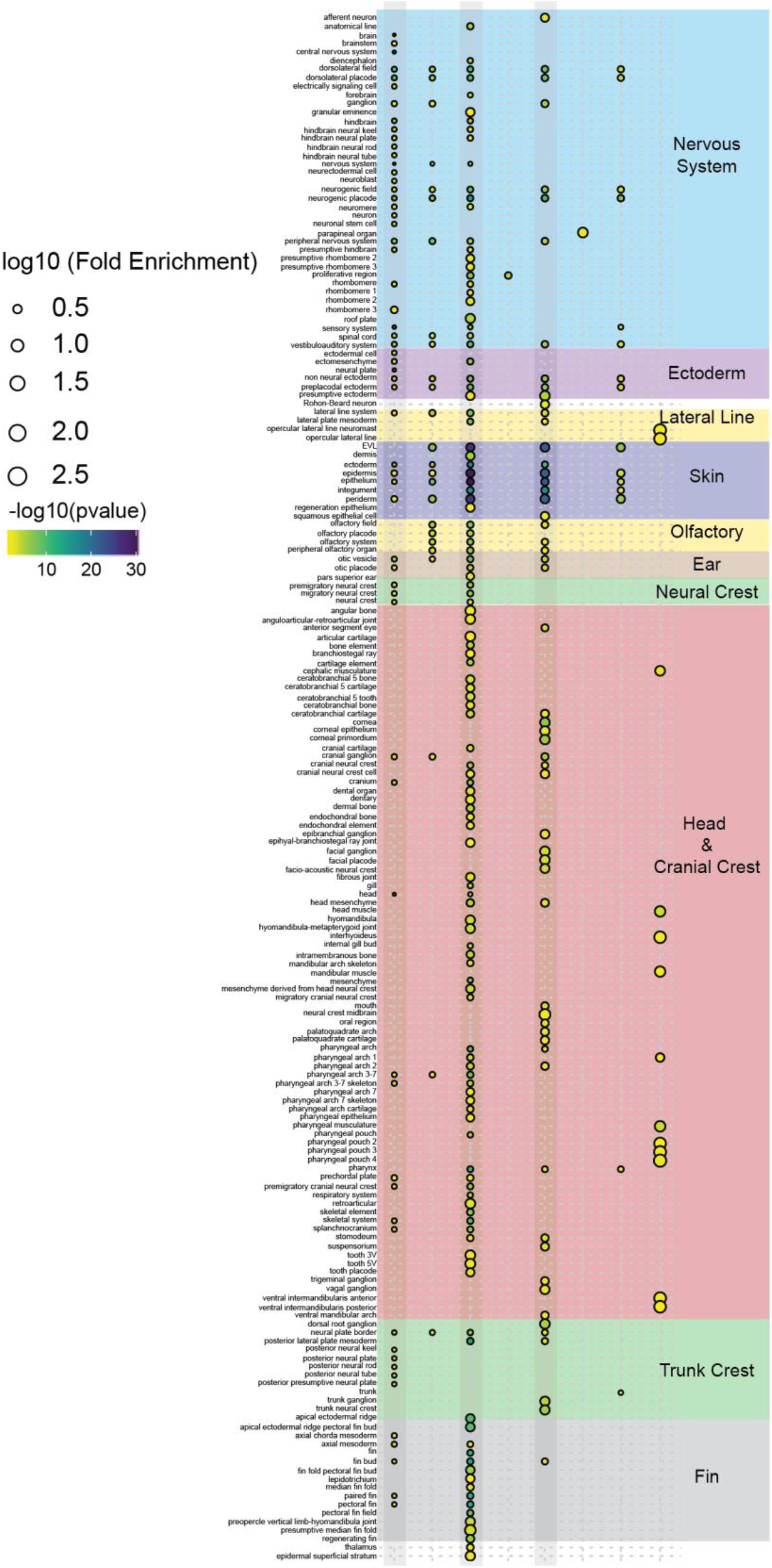
Enlargement of zebrafish anatomy enrichment from Figure 4h.

**Supplementary Figure 4.**
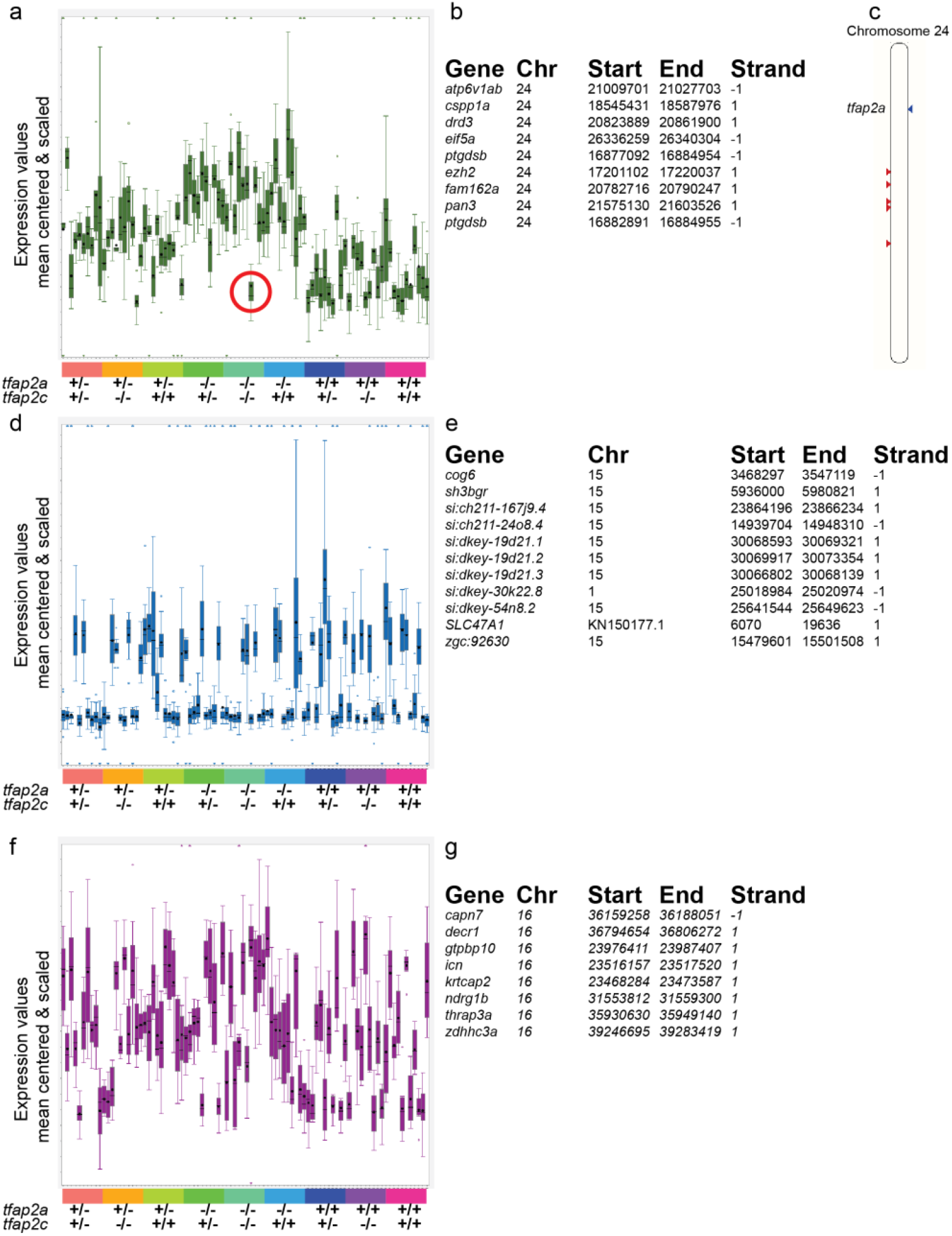
Examples of haplotype specific signals from the *tfap2a*;*tfap2c* 15 somite RNA-Seq data set. **a**, **d**, **f** Markov clusters from BioLayout3D expression analysis shown in Figure 5 a’ which contain genes linked to a specific region on a particular chromosome. A bar indicating the genotypes of the embryos is at the bottom. **a** A cluster of genes located on chromosome 24 linked to *tfap2a*. Genes behave in three groups depending on whether *tfap2a* is heterozygous, homozygous or wild-type. A recombination has occurred in one embryo in the *tfap2a* homozygous group and that cluster of genes now behaves as the wild-type condition. **b** A list of genes and their chromosomal positions which make up the cluster in **a**. **c** A karyotype map of chromosome 24 showing the location of *tfap2a* (blue arrow head right) and the positions of the genes contained in the cluster (red arrow heads left). **d** A cluster of genes on chromosome 15 where genes fall into two different groups, indicating one of the parents would have been heterozygous for the region. **e** A list of the genes contained in the cluster which are mostly on chromosome 15 and potentially two incorrectly mapped genes. **f- g** A third example of a haplotype specific region located on chromosome 16 where both parents are presumably heterozygous for the region leading to three different groups.

**Supplementary Figure 5.**
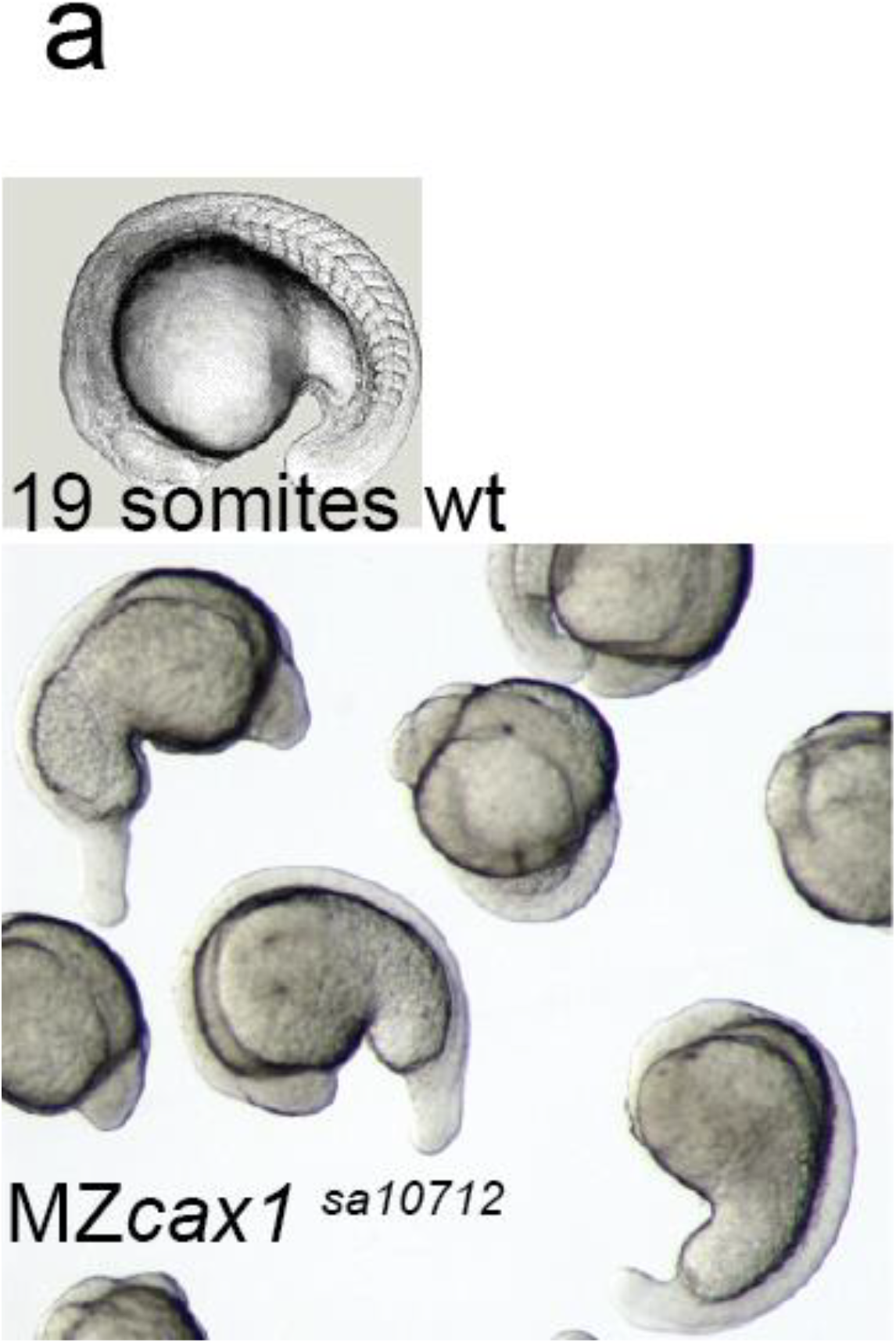
MZ*cax1^sa10712^* phenotype at 19 somites stage.

